# Identification of CD317-Positive Pro-inflammatory Immune Stromal Cells in Human Mesenchymal Stromal Cell Preparations

**DOI:** 10.1101/2022.02.10.479972

**Authors:** Alasdair G Kay, James M Fox, James Hewitson, Andrew Stone, Sophie Robertson, Sally James, Xiao-nong Wang, Elizabeth Kapasa, Xuebin Yang, Paul G Genever

**Affiliations:** York Biomedical Research Institute and Department of Biology, University of York, York, YO10 5DD, UK; Boston Children’s Hospital / Harvard Medical School, Boston MA, USA; Translational and Clinical Research Institute, Newcastle University, Newcastle, NE2 4HH; Department of Oral Biology, School of Dentistry, University of Leeds, St James’s University Hospital, Leeds, LS9 7TF

**Keywords:** Mesenchymal stromal cells, MSC subtypes, heterogeneity, immunomodulation, CD317, BST2, tetherin

## Abstract

Mesenchymal stromal cell (MSC) heterogeneity clouds biological understanding and hampers their clinical development. In MSC cultures most commonly used in research and therapy, we have identified an MSC subtype characterised by CD317 expression (CD317^pos^ (29.77±3.00% of the total MSC population), comprising CD317^dim^ (28.10±4.60%) and CD317^bright^ (1.67±0.58%) MSCs) and a constitutive interferon signature linked to human disease. We demonstrate that CD317^pos^ MSCs induced cutaneous tissue damage when applied a skin explant model of inflammation, whereas CD317^neg^ MSCs had no effect. Only CD317^neg^ MSCs were able to suppress proliferative cycles of activated human T cells *in vitro*, whilst CD317^pos^ MSCs increased polarisation towards pro-inflammatory Th1 cells and CD317^neg^ cell lines did not. Using an *in vivo* peritonitis model, we found that CD317^neg^ and CD317^pos^ MSCs suppressed leukocyte recruitment but only CD317^neg^ MSCs suppressed macrophage numbers. Using MSC-loaded scaffolds implanted subcutaneously in immunocompromised mice we were able to observe tissue generation and blood vessel formation with CD317^neg^ MSC lines, but not CD317^pos^ MSC lines. Our evidence is consistent with the identification of an immune stromal cell, which is likely to contribute to specific physiological and pathological functions and influence clinical outcome of therapeutic MSCs.

## Introduction

Mesenchymal stromal cells (MSCs, frequently referred to as “mesenchymal stem cells”) exist in bone marrow at a frequency of approximately 0.001-0.01%^1^ and are typically self-renewing for 10-50 population doublings^2,3^. MSCs can differentiate into skeletal lineages (osteogenic, adipogenic, chondrogenic) and regulate immune cell function^4^ predominantly through the release of cytokines and other immunosuppressive factors^5^. The International Society for Cell & Gene Therapy (ISCT) guidelines identifies MSCs as cells that exhibit tri-lineage differentiation *in vitro* and plastic adherence, alongside an expression profile of selected cell surface epitopes (e.g. typically presence of CD105, CD73 and CD90, and absence of CD45, CD34, CD14 or CD11b, CD79alpha or CD19 and HLA-DR)^6^. There has been some progress in identifying *in vivo* markers of MSC populations in mouse and human systems, which include LEPR, nestin, CD271, CD146 and CD164^7^, however, no single marker for MSCs exists in general use. Cells labelled as “MSCs” are used internationally in clinical trials but are rarely characterised (using ISCT or any other criteria^8^) and delivery variable success^9^.

The majority of trials assessing efficacy of MSCs currently aim to harness immunomodulatory properties^10^, though widespread clinical translation is greatly hindered by insufficient data demonstrating strong and consistent clinical effect, mechanisms of action and diverse application of selection criteria^11^. In addition, MSCs from different origins have been applied in clinical trials with varied outcomes for disorders including osteoarthritis^12-15^, osteoporotic fracture repair^16^, rheumatoid arthritis^17-19^, type 1 diabetes mellitus^20^, diabetic kidney disease^21^, multiple sclerosis^22,23^, liver failure^24-26^, amyotrophic lateral sclerosis^27-30^ and COVID-19^31-33^. Notably, although serious adverse events are extremely rare, mild, transient or acute adverse events occurring are often related to acute inflammation^13-16,19,21,25,29,30^, fever (pyrexia)^17,19,22,24,26,30,34^, infection^12,16,21,23,30^, allergic reactions/hypersensitivity^13,15,16,19^ and haematoma^13^, all of which are implicated in immune responses.

Studies examining heterogeneity in MSCs have identified multiple subpopulations of MSCs with varied potency for both differentiation and immunomodulation^35-40^. Heterogeneous populations of MSC-like cells have been isolated from both adult and neonatal sources (e.g. bone marrow^41,42^, peripheral blood^43^, adipose tissue^44,45^, synovial membrane and fluid^46,47^, dental pulp^48^, endometrium^49^, periodontal ligament^50^, tendon^51^, trabecular bone^52^, umbilical cord^53,54^, umbilical cord blood^55,56^, placenta^57^). There are further indications that MSC-like cells may be present in most vascularised tissues in some form^58,59^. This widespread distribution of MSC-like cells with varied differentiation capacities and fluctuations in the expression levels of characterising surface markers has prompted increasing reports of unipotent tissue-specific MSCs, yet bone marrow-derived MSCs are generally considered to be a population composed entirely of cells possessing tripotent differentiation capacity^6^. This raises the hypothesis that heterogeneous cell populations may collectively characterise as MSCs using ISCT (and other) criteria but comprise subsets of cells specialised to perform different functions. The widespread reporting of immunomodulatory capacities of MSCs and the impact of immune responses during tissue formation and comorbidity in degenerative disease highlights the likelihood of a nascent, endogenous population of cells that operate primarily to convey or control immune function. This population has the potential to support tissue regeneration rather than contributing to it.

We previously demonstrated the heterogeneity of human MSCs through the identification of multiple subpopulations using a clonal isolation and immortalisation strategy that enabled in-depth and reproducible characterisation^60^. These populations included an immune-primed MSC subtype identifiable through positive expression of CD317 (bone marrow stromal antigen-2 (BST2) or tetherin) and possessing enhanced immunomodulatory capacity. Here, we tested the hypothesis that CD317 positive (CD317^pos^) stromal cells function primarily to direct the immune response and do not contribute to tissue generation or repair in both physiological and pathological processes and therefore represent an identifiable MSC subtype.

## Results

### MSC identity of CD317-expressing stromal cells

In our previous work we isolated nullipotent, CD317^pos^ MSC lines (Y102 and Y202) alongside differentiation-competent, CD317^neg^ MSC lines (Y101 and Y201) from the same heterogeneous donor source suggesting that a subpopulation of stromal cells exists in typical MSC preparations but may not contribute to ‘classic’ MSC functions. Here, we examined the stromal phenotype the CD317^pos^ and CD317^neg^ MSC lines. An *in silico* assessment using the Rohart Test^61^ was applied to accurately discriminate MSCs from fibroblasts, other adult stem/progenitor cell types and differentiated stromal cells. This test uses 16 key MSC marker genes as a proven panel of identifiers that has independently confirmed MSC status with 97.85% accuracy in 635 cell samples^61^. All of the immortalised CD317^neg^ and CD317^pos^ stromal cell lines maintained gene expression patterns that independently confirmed their MSC status (Figure S1A and Table S1).

Next, we used mass spectrometry to determine cell surface protein expression profiles across the different cell lines. We identified a high number of commonly expressed proteins alongside cell line-specific variations. Using a false detection rate of 3%, we found 2678 proteins expressed across all MSC lines, with 584 (65.2%) of these commonly expressed (Figure S1B), which may reveal a common stromal surfaceome signature. Percentage similarity at the surfaceomic level ranged from 76.0% to 83.5% (Figure S1C). Unique proteins were identified in Y101 (20 proteins, 2.2%); Y102 (30 proteins, 3.3%); Y201 (36 proteins, 4.0%); and Y202 (21 proteins, 2.3%). These analyses also confirmed that CD317 (BST2) was only identified on Y102 and Y202 MSC lines. Principle component analysis (PCA) was used to aid interpretation of mass spectrometry data through dimensionality reduction. Analysis highlighted that MSC lines clustered distinctly within the whole population but were on a similar spectrum of observation, with Y102 and Y202 lines lying further from the mean of the whole population (Figure S1D). Together, these data demonstrate that the CD317^neg^ Y101 and Y201 cell lines, and the CD317^pos^ Y102 and Y202 cell lines have broadly similar protein expression profiles in common with other MSC preparations and may be used as models for different MSC subtypes.

### Identification of CD317^dim^ and CD317^bright^ populations in primary MSCs

We previously reported a CD317^pos^ MSC subset with average frequency of 1-3% in low passage primary MSCs^60^. Here, using flow cytometry analysis with Y201 and Y202 populations gating for primary cells as either CD317^neg^ or CD317^pos^, we were able to demonstrate that CD317 positivity can be subdivided into CD317^dim^ and CD317^bright^ populations in primary MSC cultures (Figure 1A, S1E). Further examination of n=24 primary MSC populations recorded proportions at CD317^neg^ (70.57±5.09%) and CD317^pos^ (29.77±3.00%), comprising CD317^dim^ (28.10±4.60%) and CD317^bright^ (1.67±0.58%) (Figure 1B). We observed a decrease in CD317 expression over time in culture (passages 1-4), however this trend did not reach statistical significance due to the variability of initial proportions of CD317^pos^ cells when CD317^dim^ was included as a CD317 positive result (means passage 1 = 50.66±27.63%, passage 2 = 30.35±6.03%, passage 3 = 26.07±11.78%, passage 4 = 22.18±12.26%; n=2,12,7,3) (Figure S1F). We made a similar observation when examining subsets of CD317^dim^ and CD317^bright^ cells, with CD317^bright^ cells almost absent by passage 4 (Figure 1C). CD317 expression in isolated primary MSCs from passage 3 to 4 reduced by 49.01 ± 11.84% (n=5); with a freeze/thaw cycle at passage 3, this reduction was recorded at 63.94 ± 3.64% in the same cells (n=5) (Figure S1G). Therefore, human primary MSC isolates express CD317 on a spectrum that varies from cell to cell and from individual to individual; the overall proportion of CD317^pos^ MSCs, as a composite of CD317^dim^ and CD317^bright^, is 28-29% in heterogeneous MSC cultures (combining all analyses of primary cell donors, percent CD317^pos^ MSCs is 28.44±3.82% (mean ± SEM), range of 0.01-93.03%; median=19.89%; n=52). Within CD317^pos^ cells, there was no difference in percentage CD317 expression based upon donor gender (mean expression female 40.02±5.27; male 24.77±6.51; Mann Whitney T-test p=0.051, n=52) or correlation between donor age and CD317 expression (mean age: 69.75±1.29 years; range 45-88; Pearson correlation p=0.141, n=52),) (Figure 1D, 1E). There was, however, a significant negative correlation between CD317 expression and BMI (mean 28.06±0.78; range 17-44; Spearman correlation p<0.05, n=52) (Figure 1F).

**Figure 1.**
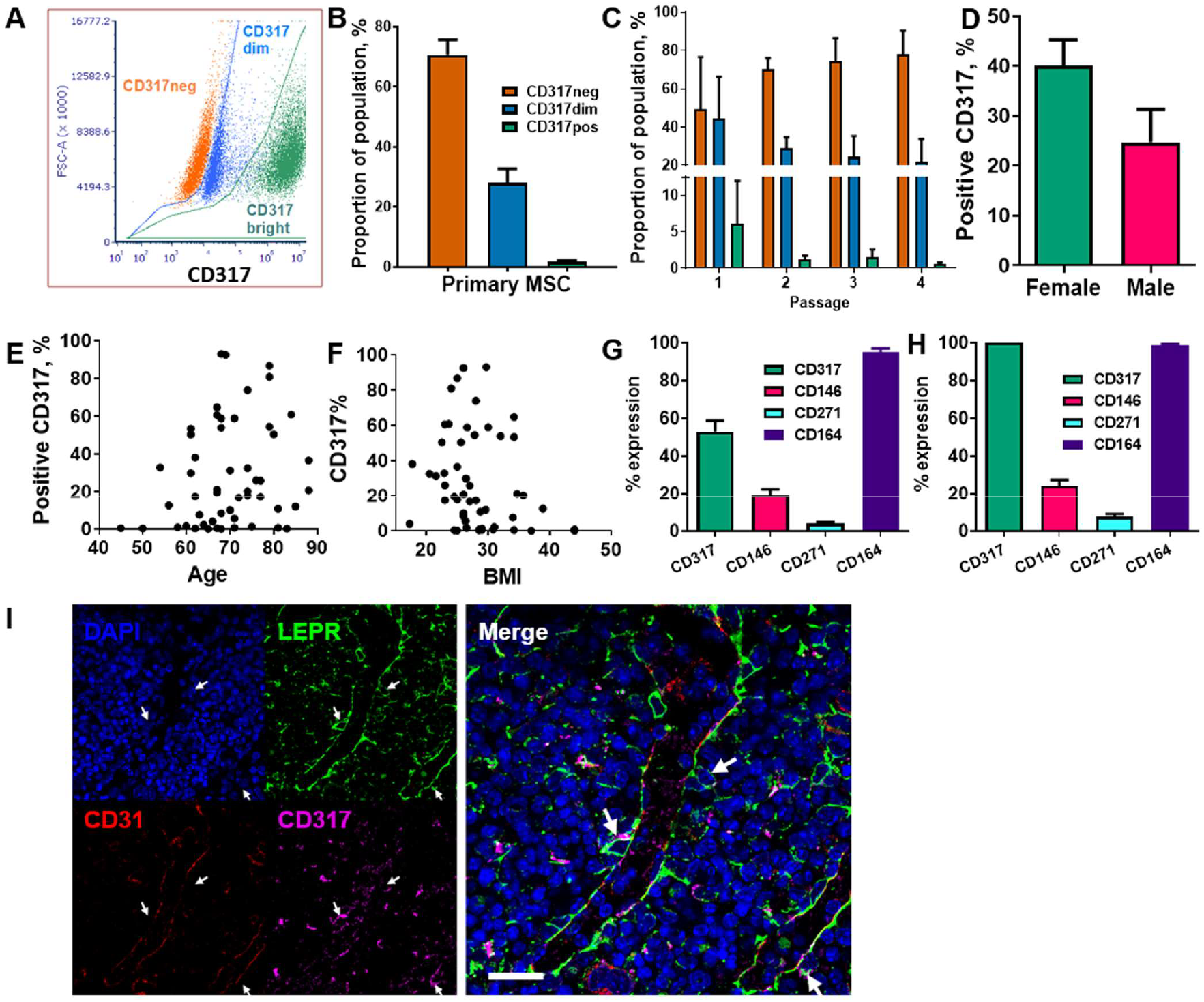
Analysis of CD317-expressing MSC populations within primary cell isolates. (A) The CD317 expressing populations can be divided into CD317^bright^ and CD317^dim^ with CD317^bright^ MSCs. (B) Average proportions of CD317^neg^ and CD317^pos^, comprising CD317^dim^ and CD317^bright^, in primary MSCs lines. (C) Expression of CD317 over early passages 1 to 4 in Primary MSCs with CD317^neg^ increasing, CD317^dim^ and CD317^bright^ decreasing during in vitro culture (n=2-12). Variation of CD317 expression with gender (D), age (E) and BMI (F) in primary donors (n=52). (G) Isolated MSCs from human primary donors showed CD317^pos^ (CD317^dim^ and CD317^bright^ combined) with mean values of CD317^pos^, CD146^pos^, CD271^pos^ and CD164^pos^ (n=27). (H) Examination of the CD317^pos^ population only, showed similar proportions of each marker to those seen in the whole population (n=27). (I) CD317 expression was detected throughout the bone marrow of mice with low frequency colocalization of CD317 and LEPR+ in peri-sinusoidal regions (arrows).

We previously demonstrated that the hTERT immortalised MSC lines display typical (ISCT) surface marker profiles^60^. Here, we also examined surface markers commonly associated with human stromal progenitor cells or subsets, including CD146, CD271 and CD164, within CD317^neg^ and CD317^pos^ primary MSC populations. Isolated MSCs from human primary donors showed CD317^pos^ (CD317^dim^ and CD317^bright^ populations combined) with mean % expression values of CD317^pos^ (52.90±5.89%), CD146^pos^ (19.46±3.07%), CD271^pos^ (4.025±0.71%) and CD164^pos^ (95.03±2.11%) (n=27) (Figure 1G). Examination of the CD317^pos^ population only showed similar proportions of each marker to those seen in the whole population: CD146^pos^ (24.21±3.23%), CD271^pos^ (7.78±1.35%) and CD164^pos^ (97.18±0.66%) (n=27) (Figure 1H). These findings demonstrate that expression of these markers is independent of CD317 positivity and that CD164 identifies virtually all CD317^neg^ and CD317^pos^ MSCs.

Comparative gene expression analysis has previously demonstrated a correlation between murine peri-sinusoidal stromal cells and CD317^pos^ MSCs^62^. LEPR has been shown to mark peri-sinusoidal stromal cells in mouse tissue^63^. Here we investigated CD317^pos^/LEPR^pos^ stromal cells in mouse bone marrow to identify the *in vivo* location of this subpopulation. CD317 expression was detected throughout the bone marrow with low frequency colocalisation of CD317 with LEPR restricted to peri-sinusoidal regions adjacent to CD31-positive endothelial cells (Figure 1I).

### Immune profile of CD317^pos^ MSCs

Our previous transcriptomic data indicated that CD317^pos^ Y102 and Y202 MSC lines display a constitutive immunostimulatory expression profile^60^, which we sought to define here using the MSC lines and primary cells sorted based on CD317 expression. We confirmed by qPCR that ICAM1 (CD54) mRNA levels were significantly elevated in CD317^pos^ Y102/Y202 compared to CD317^neg^ Y101 (Figure 2A). Although ICAM1 mRNA expression levels appeared similar in primary MSCs sorted for CD317 positivity (Figure 2A), flow cytometric analysis demonstrated that cell surface ICAM1 expression, as shown by mean fluorescence intensity (MFI), was significantly increased on CD317^pos^ primary MSCs versus CD317^neg^ MSCs and CD317^pos^ Y102/Y202 versus CD317^neg^ Y101/Y201 (Figure 2B). Comparative analysis of CXCL10 and CXCL11 mRNA levels in immortalised MSC lines and primary MSCs sorted for CD317 demonstrated significantly increased expression in all CD317-positive MSCs compared to CD317-negative counterparts (n=7; experiments performed in triplicate) (Figure 2C, 2D).

**Figure 2.**
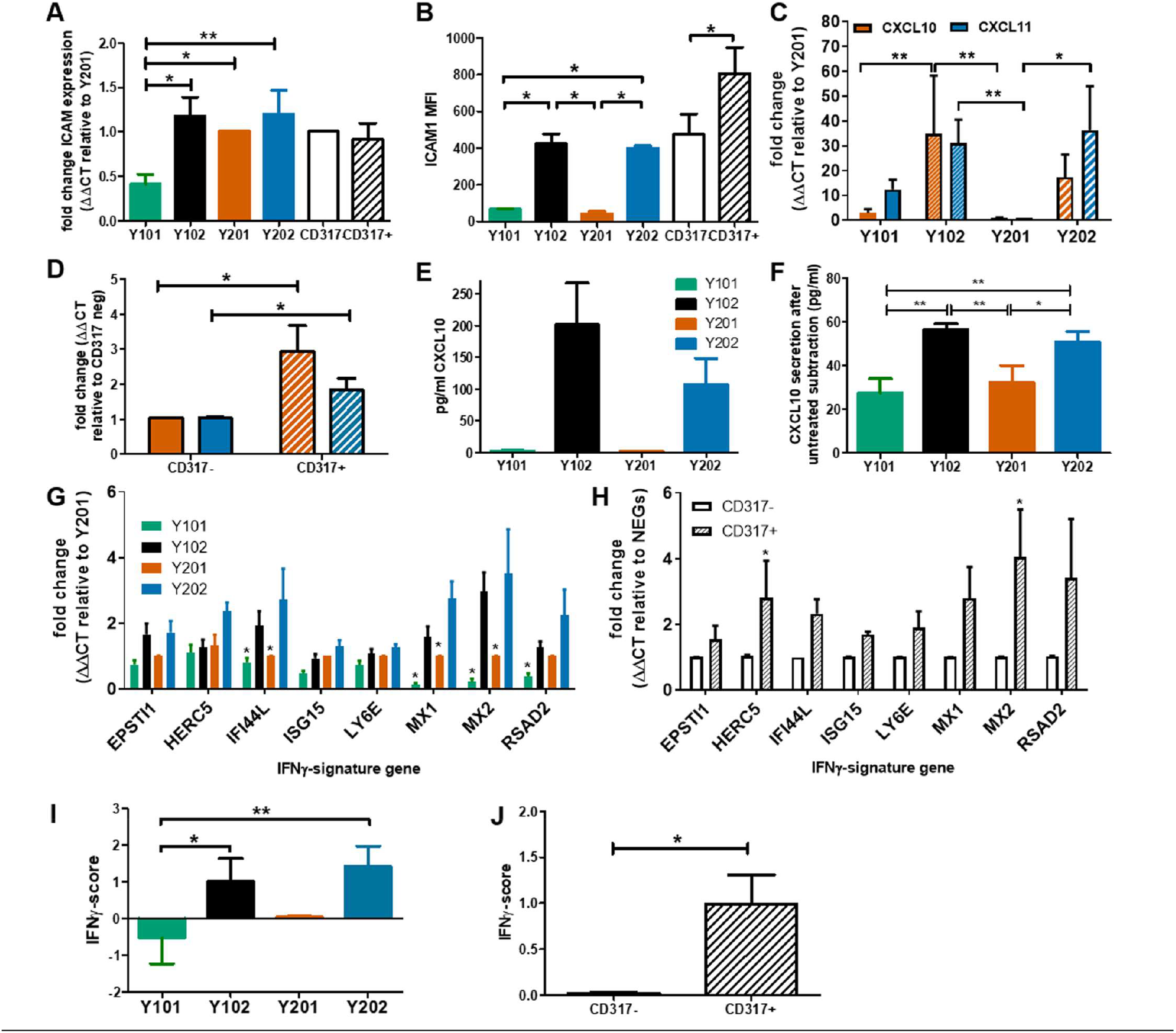
Examination of the immune profile of CD317pos MSCs. (A) Comparative mRNA expression of ICAM-1 in MSC lines and primary cells sorted by CD317 expression (RNA was extracted from 3 different donors or 5 cell line passages; qPCR performed in triplicate, mean shown ± SEM). (B) Mean fluorescence intensity of ICAM-1 expression on the cell surface of MSC lines and primary MSCs differentially gated by CD317 staining (MSCs from 5 different donors or 4 different passages of MSC lines were stained for flow cytometry, mean shown ± SEM). (C)/(D) Comparative (mean ± SEM) mRNA expression of CXCL10 (red) and CXCL11 (blue) in MSC lines/ primary MSCs sorted for CD317 expression (RNA was extracted from 7 different donors/7 different cell passages; experiments were performed in triplicate). (E/F) CXCL10 secretion by MSC lines prior to IFN-γ priming and after priming with baseline (unprimed) secretion subtracted (mean ± SEM, n=2). (G/H) Comparative mRNA expression of 8 IFN-γ signature genes in MSC lines/primary MSCs sorted by CD317 expression (RNA was extracted from 5 different donors/5 different cell passages; experiments were performed in triplicate, mean shown ± SEM). (I)/(J) IFN-γ score for MSC lines/primary MSCs sorted by CD317 expression (n=5)*/** = significance at P<0.05/0.01 using an appropriate statistical test.

CD317, ICAM-1 and CXCL10 are regulated by interferon-gamma (IFN-γ). We analysed expression levels of the IFN-γ receptor by flow cytometry and demonstrated that it was expressed at similar levels in all four MSC lines, independent of CD317 expression (MFI, Y101=9.11, Y201=8.41, Y102=9.60, Y202=9.84; p>0.05) (Figure S2A). This finding suggested that all MSC lines were capable of responding to IFN-γ stimulation in a similar manner, but CD317-positive MSCs may be primed to transduce IFN-γ stimulation more effectively. Secretion of CXCL10 was measured with (Figure 2E) and without (Figure 2F) IFN-γ exposure. Under basal, unstimulated conditions, CD317^pos^ Y102/Y202 MSCs secrete larger amounts of CXCL10 compared to CD317^neg^ Y101/Y201. Following IFN-γ priming, CD317^pos^ MSC lines demonstrate a significantly increased ability to secrete additional amounts of CXCL10 compared to CD317^neg^ MSC lines. However, IFN-γ has a proportionally much larger stimulatory effect on CXCL10 secretion by CD317^neg^ Y101/Y201 cells, suggesting that constitutive interferon signalling is a feature of CD317^pos^ MSC lines (Figure 2F).

Examination of a further panel of eight IFN-γ related genes showed remarkably different expression between CD317^pos^ and CD317^neg^ MSCs (Figure 2G, 2H). Using a method described by Raterman *et al*^64^, we generated an IFN-γ signature score for CD317^pos^ and CD317^neg^ MSCs using the average of the log base-2 normalised relative fold changes of the eight IFN-γ related genes. We demonstrated that CD317^pos^ MSC lines and primary MSCs had a significantly increased IFN-γ signature score compared to CD317^neg^ MSCs (Figure 2I & 2J).

Bioinformatics analysis of differentially expressed genes (DEGs) using combined transcriptomic data^60^ from CD317^neg^ (Y101 & Y201) and CD317^pos^ (Y102 & Y202) MSC lines identified 2340 significantly upregulated genes in CD317^pos^ MSC samples (FC>2, p<0.05) with clear clustering of the Y01 group (Y101, Y201) and the Y02 group (Y102, Y202) (Figure S2B). The 10 most significantly upregulated genes in the CD317^pos^ group were immune-related and/or interferon-regulated, including OAS1, OASL, RSAD2 and CD317 (BST2) (Figure S2C). IFN signalling and elevated IFN-signatures are associated with different human disease states^65^. When comparing the upregulated Y102/Y202 gene sets with six publicly available transcriptomic databases for autoimmune and related disorders (Table S2), we identified a significant association between DEGs and GO terms that were enriched in Y102/Y202 MSC lines and psoriasis, eczema and, to a lesser extent, rheumatoid arthritis and osteoporosis (Table S3). Similar observations were made when comparing enriched signalling pathways across Y102/Y202 and disease datasets (Table S4).

Therefore, a resident MSC subtype can be identified as CD317^pos^ICAM-1^hi^CXCL10^hi^ with apparent constitutive interferon signalling, which is likely to contribute to specific physiological and pathological immune functions.

### Roles of CD317^pos^ and CD317^neg^ MSCs in monocyte and T cell function

Immunomodulation may be affected through paracrine signalling altering cell recruitment and retention in response to signalling molecule expression. The CCL2 receptor, CCR2, is a monocyte chemoattractant receptor protein involved in macrophage activation in cells expressing high levels of CCL2. Significantly higher CCL2 mRNA expression and protein secretion was detected in CD317 expressing MSCs versus CD317-negatives (Figure 3A & B).

**Figure 3.**
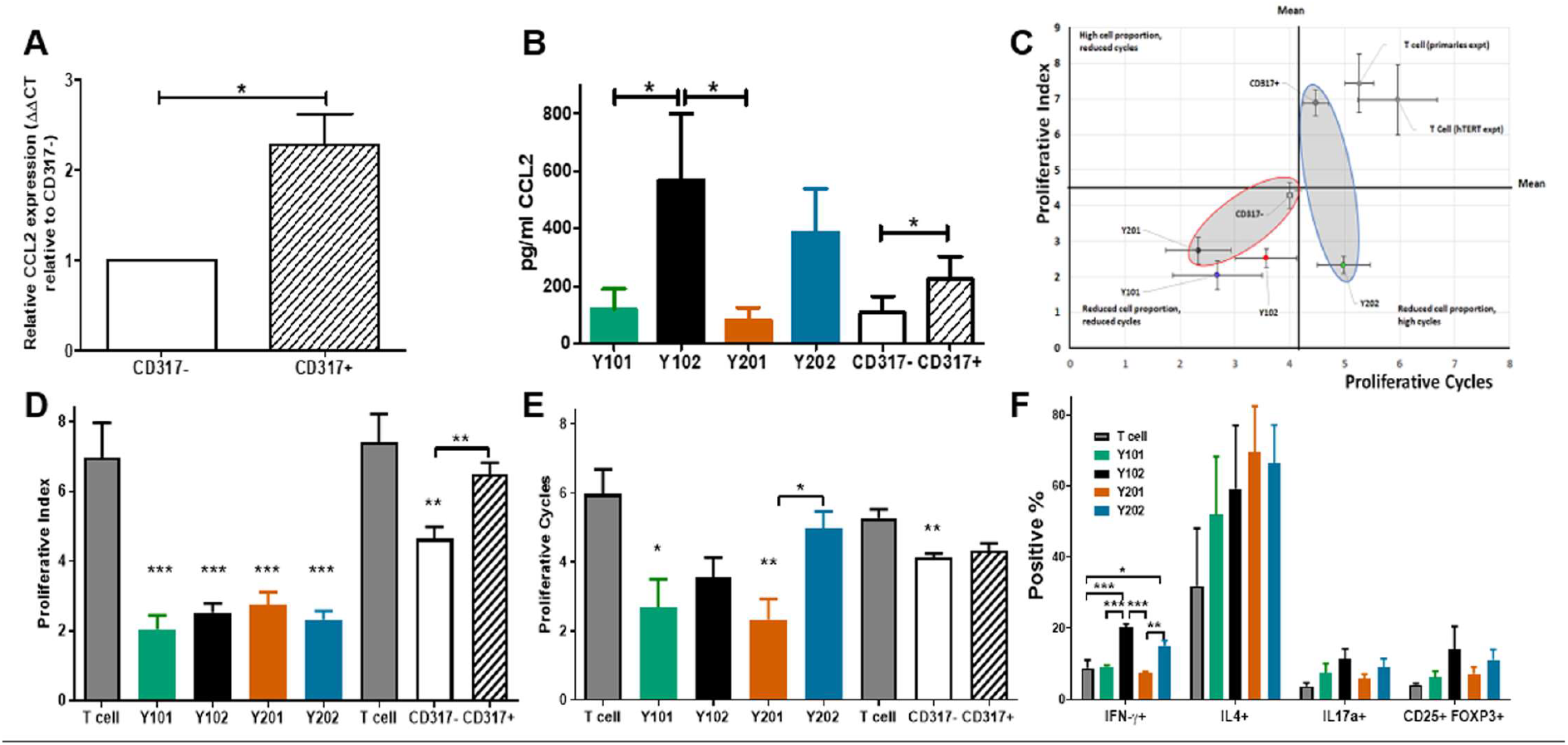
Influence of CD317^neg^ MSCs and of CD317^pos^ MSCs on immune cell function. (A) Comparative mRNA expression of CCL2 in primary MSCs sorted by CD317 expression (RNA was extracted from 7 different donors; experiments performed in triplicate, mean shown ± SEM). (B) CCL2 secretion in primary MSCs sorted by CD317 expression and MSC lines (from 4 different donors/4 different cell line passages; experiments performed in triplicate, mean shown ± SEM). (C) In vitro co-culture of hTERT immortalised lines Y201 and Y202 and primary CD317^neg^ and CD317^pos^ cells with activated T cells. CD317^neg^ cells reduce proportion of proliferating T cells and number of cell cycles achieved (D) hTERT cell lines significantly reduce proportion of proliferating cells as demonstrated through proliferative index (E) CD317^neg^ cell lines reduce proliferative cycles achieved by activated T cells in comparison to CD317^pos^ or T cell alone controls. (F) assessment of the influence of MSC on T cell polarisation in co-culture demonstrates CD317^pos^ cells influence activated T cells to preferentially polarise towards IFN-γ expressing (Th1) subset with indications of increased IL17a+ and CD25+FOXP3+ expressing cells.

In the presence of an antagonist for CCR2, migration of monocytic cells (THP-1) towards supernatant from CD317-expressing MSC lines was selectively inhibited compared to CD317-negative MSC lines (Y101, Y201 vs Y102, Y202; 19.37±9.57, 19.61±8.89 vs 39.01±6.57, 41.02±4.79) (Figure S3A). We tested whether the supernatant of CD317^pos^ and CD317^neg^ MSCs could induce the migration of both monocytic (THP-1) and T cell (HUT-78) lines in transwell assays. We demonstrated that both THP-1 and HUT-78 cells migrated towards MSC supernatants suggesting that MSCs secrete both monocyte and T cell chemoattractants (Figure S3B).

MSCs have previously been shown to suppress activated T cell proliferation whilst maintaining inactivated T cell viability in co-culture^66^. Several mechanisms are proposed that provide evidence for IFN-γ mediated immunosuppression^67^, potentially achieving MSC deactivation of T cells through IFN-γ receptor targeting or IFN-γ-mediated induction of indoleamine 2,3-dioxygenase (IDO) from MSCs, whereby tryptophan is catabolised leading to suppression of T cell proliferation and subsequent apoptosis of activated T cells, leaving inactivated T cells in a viable state^68,69^. In this work, T cell proliferation was assessed for peaks of gradual division (proliferative index)^70^ and proliferative cycles (population doublings)^71^ over 5 days of co-culture with or without CD317^pos^ and CD317^neg^ MSC cell lines (Figure S3C). T cells do not proliferate in culture, unless activated with anti-CD3/CD28, and undergo cell death in absence of IL-2, which is produced *in vivo* by activated T cells^72^. Compared to T cells alone, all MSC lines and CD317^neg^ primary MSCs significantly reduced proliferative index scores, whereas CD317^pos^ primary MSCs had no significant effect on T cell proliferative index (Figure 3C, 3D). Assessment of T cell proliferative cycles showed significant reductions when cultured with CD317^neg^ Y101/Y201 and CD317^neg^ primary MSCs (Figure 3C, 3E) compared to T cells alone. However, CD317^pos^ Y102/Y202 MSCs and CD317^pos^ primary MSCs did not significantly reduce the number of proliferative cycles, although a decline was observed (Figure 3C, 3E). These results demonstrate that CD317^pos^ MSCs are capable of inactivating a proportion of proliferating T cells, although this effect is not sufficient to reduce the number of proliferative cycles that the residual activated cells achieve, pointing to a diminished immunosuppressive function for CD317^pos^ MSCs.

Next, we determined the effect of CD317^neg^ and CD317^pos^ MSCs on the polarisation of naïve T cells into effector lineages with immunosuppressive/anti-inflammatory function. CD317^pos^ MSC lines induced a significant increase in the development of pro-inflammatory Th1 cells. Both Y102 (20.32 ± 0.92%, p<0.001) and Y202 (15.11 ± 1.46%, p<0.05) increased Th1 polarisation, as indicated by IFN-γ expression, in comparison to T cells alone (8.79±2.30%), CD317^neg^ Y101 (9.25±0.42%, p < 0.001 (Y102)) and Y201 (7.31±0.60%, p <0.001 (Y102), p <0.01 (Y202)) (One way ANOVA with Bonferroni post hoc test). An increase was also observed in Th2 cells for all MSC lines (p>0.05, n.s.). Both Th17 and Treg cells, as indicated by IL17a and CD25/FOXP3 expression respectively, increased slightly with CD317^pos^ MSC lines, but not statistically significantly. By examining total proportions of differentiating cells, it was notable that a large proportion of CD4+ T cells cultured alone did not commit to any lineage when compared to co-culture with MSC lines. When proportions are summated, only 48.49% of T cells cultured alone differentiated into the 4 lineages examined, whilst approximately 75% (Y101), 90% (Y201) and 100% (Y102, Y202) differentiation into these lineages was observed when T cells were co-cultured with MSC lines (Figure 3F).

### Pro-inflammatory and Immuno-regulatory potential of CD317^neg^ and CD317^pos^ MSCs in vitro and in vivo

Considering the stark differences in immune profiles of CD317^neg^ and CD317^pos^ MSCs, we tested their effects in different inflammatory models. Prior to *in vitro* and *in vivo* testing, we confirmed the representative CD317^neg^ and CD317^pos^ MSCs (Y201, Y202) were not affected by viral contamination as a potential origin or contributor to constitutive IFN-γ expression. All cell samples were tested in triplicate and returned negative results for molecular diagnostics of infectious diseases (Human Comprehensive CLEAR Panel, Charles River) using PCR for RNA representing a panel of 26 virions.

Initially, we investigated the potential pro-inflammatory property of CD317^neg^ Y201 and CD317^pos^ Y202 MSCs in a skin explant model, which is an *in vitro* tool to detect the presence of cutaneous tissue damage following a pro-inflammatory insult^73,74^. CD317^neg^ Y201 and CD317^pos^ Y202 MSCs were primed with IFN-γ or TNF-α and co-cultured *in vitro* with skin explants.

In this assessment, no tissue damage was observed after skin co-incubation with CD317^neg^ Y201 cells in all conditions tested (Figure 4A top panel and Figure 4B left panel). In contrast, cutaneous tissue damage was detected when skin was co-cultured with unstimulated or TNF-α stimulated CD317^pos^ Y202 cells showing clear cleft formation in the basal layer between the dermis and epidermis (Figure 4A bottom panel and Figure 4B right panel). When comparing the ability to cause tissue damage, Y202 cells caused significantly increased damage compared to Y201 cells in unstimulated and TNF-α stimulated conditions (p<0.05) whilst no cutaneous tissue damage was observed when skin was co-cultured with IFN-γ stimulated Y202 cells.

**Figure 4.**
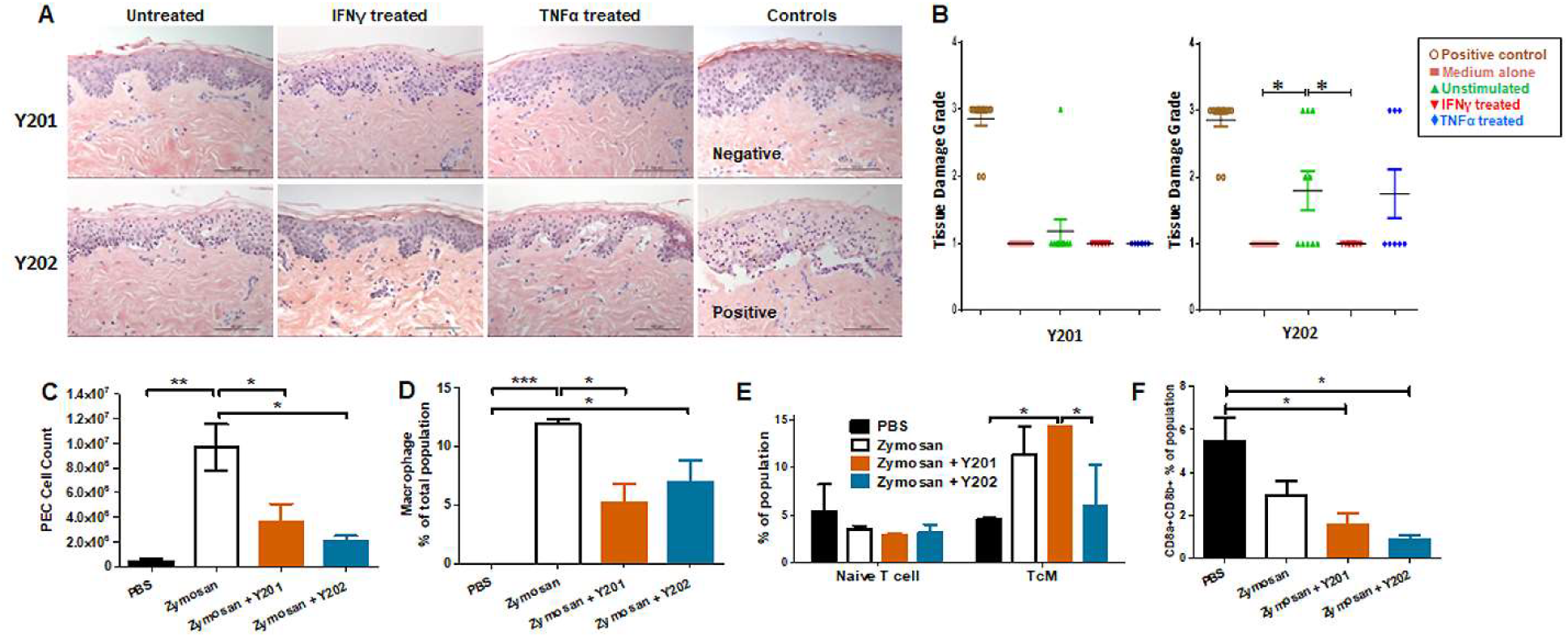
In vitro and in vivo immunomodulation by CD317^neg^ Y201 or CD317^pos^ Y202 MSCs. (A) Representative images of skin explants independently assessed for damage to tissues, examining keratinocytes, basal cells, keratotic bodies, the appearance of sub-epidermal clefts at the junction with the dermis and in highly damaged tissue the appearance of complete epidermal separation following treatment with MSCs primed with IFN-γ or TNF-α and co-cultured *in vitro*. (B) Y201 co-culture did not prompt damage to the tissue in any conditions whilst Y202 cell line demonstrated marked tissue damage in untreated cells and TNF-α treated cell lines. Both Y201 and Y202 cell lines retained the ability to inhibit tissue damage when primed with IFN-γ. (C) MSCs subsequently applied to an *in vivo* peritonitis model of inflammation showed immunomodulation through reduced immune cell recruitment, (D) reduced macrophage development following Y201 treatment, (E) increased central memory T cell development following Y201 treatment and (F) reduced CD8+ cytotoxic T cell development following Y202 treatment. n=3, *p<0.05, **p<0.01, ***p<0.001

Interferon signalling genes are regulated by interferon in host-pathogen interactions. It is hypothesised that constitutive interferon signalling occurs to provide a rapid response to pathogen infections through pre-established interferon signature^65^, such as that observed here in CD317^pos^ MSCs. To investigate the potential for constitutive IFN-γ related signalling on innate immune responses *in vivo*, we evaluated immune regulation by CD317^neg^ and CD317^pos^ MSCs in a zymosan-induced peritonitis model of acute inflammation that promotes the recruitment of monocytes and neutrophils to the peritoneal cavity. Following zymosan treatment, peritoneal exudate cells (PEC) were collected by lavage and analysis performed on the cell content. A gating strategy was devised for flow cytometric analysis of multiple PEC cell types focusing on haematopoietic, myeloid and lymphoid cells including monocytes, macrophages and T cells (Figure S4A & S4B). Treatment with either Y201 or Y202 MSC lines suppressed the recruitment of inflammation-related cells to the area. There was a significant reduction in total cells recruited in both Y201 (3.552±1.543 × 10^6^) and Y202 (2.076±0.421 × 10^6^) treated conditions compared to zymosan-induced peritonitis without treatment (9.686±1.894 × 10^6^) (p<0.05), with no significant difference between MSC-treated animal PEC numbers and PBS controls (4.420±1.790 × 10^5^) (Figure 4C).

Examination of the composition of PEC showed that zymosan-induced peritonitis prompted a significant increase in haematopoietic cells (p<0.05). No difference in recruitment of eosinophils or neutrophils was observed in MSC-treated mice when compared to zymosan alone or PBS controls (Figure S4C & S4D). Examination of the production of monocytes and macrophages in PEC samples showed no differences in monocyte recruitment, however both zymosan alone and zymosan plus Y202 showed significant increases in macrophage proportions compared to PBS controls (p<0.001, p<0.05 respectively) whilst Y201 treatment suppressed macrophage numbers (p<0.05) (Figure 4D). Within these monocyte and macrophage populations, the proportions of Ly6C positive and negative cells matched the proportions seen in zymosan treatment only animals (Figure S4F & S4G). Ly6C positive monocytes and macrophages are linked with pro-inflammatory responses by CCR2/CCL2 mediated homing to sites of tissue injury, whilst Ly6C low or negative monocytes and macrophages are reparative, guided by VCAM-1 and other adhesion proteins^75,76^.

Spleens retrieved from MSC-treated and control mice were homogenised and analysed for naïve and polarised T cells, and memory T cells. No differences were found in the mass or cellularity of spleens between controls and MSC-treated animals (data not shown). When tested, a significant increase was found in activated CD4+ central memory T cells (TcM) in CD317^neg^ Y201 cell treated conditions (14.23±0.06%) in comparison to PBS controls (4.53±0.18%) or Y202 treated animals (5.89±4.30) (Figure 4E). CD4+ effector T cell polarisation was not altered by introduction of zymosan or MSC treatments within the 24 hour time period measured. However, treatment with either CD317^neg^ Y201 (1.51 ± 0.57%) or CD317^pos^ Y202 (0.84 ± 0.25%) MSCs suppressed CD8a/b+ expression representative of cytotoxic T cell production in mice in comparison to CD8a/b+ expression in untreated animals (5.42 ± 1.10%) (Figure 4F).

### In vivo tissue formation is enhanced in CD317^neg^ MSC lines when compared to CD317^pos^ subpopulations

We hypothesised that the immunomodulatory enhancements observed in CD317-positive MSCs would impact on their tissue-forming capacity. To test this hypothesis, CD317^neg^ (Y201) and CD317^pos^ (Y202) MSC lines were loaded onto hydroxyapatite (HA) scaffolds and implanted subcutaneously in nude mice. Scaffolds were retrieved at 3 and 8 weeks post-implantation and examined using histological analysis for *de novo* tissue formation by deposition of extracellular matrix (ECM), collagen and neoangiogenesis.

CD317^neg^ Y201 MSCs showed clearly advanced ECM and collagen deposition in histological stains using Sirius Red for collagen formation and Alcian Blue for proteoglycan synthesis (Figure 5A, 5B & 5C), suggestive of a more stable capacity for tissue formation. Haematoxylin and eosin staining showed evidence of tissue formation from 3 weeks post implantation in CD317^neg^ MSCs alongside evidence at 8 week timepoints of capillary tube structures containing blood cells indicative of neoangiogenesis (Figure 5D). Although there was some evidence of tissue formation in CD317^pos^ Y202-loaded scaffolds, the tissue formed was less continuous or cohesive compared to CD317^neg^ Y201 samples and by 8 weeks post-implantation there was clear evidence of disaggregation and cleft formation at the surface of HA particle clusters following histological staining for ECM formation (Alcian Blue and Sirius Red) with no detectable vessel formation (Figure 5A, 5B, 5C & 5D).

**Figure 5.**
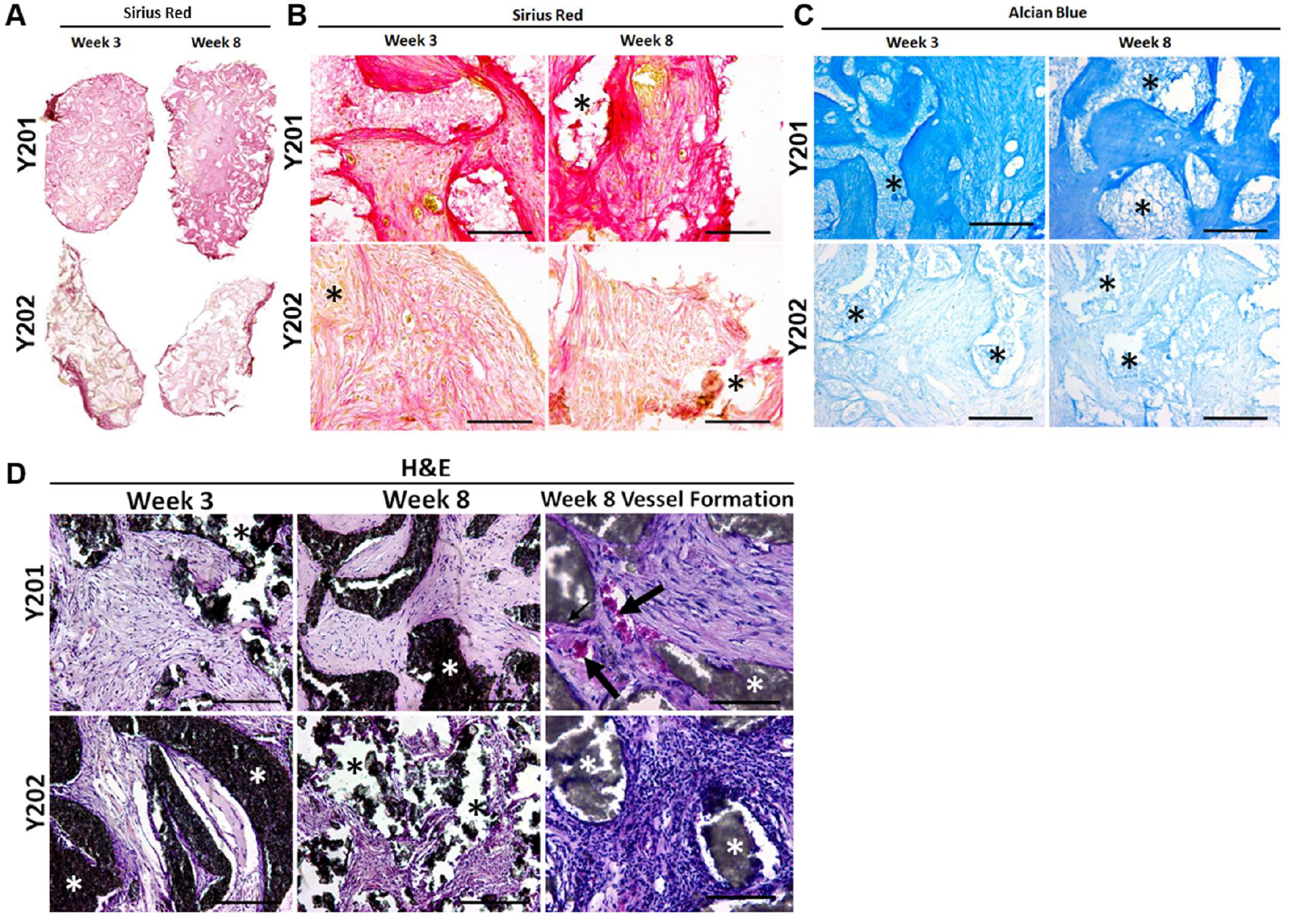
In vivo tissue generation in HA scaffolds loaded with CD317^neg^ Y201 or CD317^pos^ Y202 MSCs. (A, B) Histological staining of recovered implants using Sirius Red for collagen formation and (C) Alcian Blue for proteoglycan synthesis at 3 and 8 weeks post-implantation in HA scaffolds loaded with either CD317^neg^ Y201 MSCs and CD317^pos^ Y202 MSCs. (D) Haematoxylin and eosin staining comparting tissue and blood vessel formation at 3 and 8 weeks post-implantation in HA scaffolds loaded with CD317^neg^ Y201 MSCs and CD317^pos^ Y202 MSCs. Scale bars = 250μm. Asterisks = HA particles, arrows = blood vessels.

## Discussion

This study investigated the characteristics and properties of a CD317^pos^ subpopulation within heterogeneous MSCs and their ability to contribute to immune responses and tissue repair. We used immortalised MSC model lines and primary MSCs isolates to elucidate the biology and potential impact on the therapeutic application of these cells. Here, we confirm CD317^pos^ MSCs represent a subpopulation of cells commonly found in human MSCs preparations with an equal distribution in a range of demographic groups and health conditions. Using *in vitro* and *in vivo* functional assays, we demonstrate that CD317^pos^ MSCs have reduced immunomodulatory and tissue-forming capacity compared to CD317^neg^ MSCs, suggesting that CD317^pos^ cells will not contribute to tissue repair or *de novo* tissue formation. Any contribution of CD317^pos^ cells in therapy, when delivered within an undefined heterogeneous MSC culture, is therefore likely to be through immunomodulatory influence, and the contribution to the regenerative process is dependent upon the therapeutic target and the inflammatory environment present in the recipient at the time of transplantation. Given the potential for CD317^pos^ MSCs to respond to the inflammatory environment *in vivo*, these cells may serve a positive function in assisting the repair of damaged tissues by CD317^neg^ MSCs when transplanted as part of a heterogeneous population. However, our *in vivo* results demonstrate that CD317^neg^ cells are capable of inducing both anti-inflammatory immunomodulation and tissue regeneration in the absence of CD317^pos^ counterparts, suggesting the support function is not vital to successful repair of damaged tissue by CD317^neg^ MSCs alone. Of note, when supplied in sufficient numbers CD317^pos^ MSCs are capable of causing tissue damage, as observed in our skin explant model, which may be linked to their pro-inflammatory characteristics.

Inflammation serves a dual role in tissue repair. Cells in the immune response, such as neutrophils, function to initiate the repair process. Neutrophils cause tissue breakdown during inflammation but in the absence of neutrophils, macrophages rapidly recruited to the site of injury will display reduced rate of tissue regeneration owing to the presence of cell debris normally phagocytosed by neutrophils^77^. Our results from MSC treatment of zymosan-induced peritonitis in mice showed a neutrophil population present in PEC suspensions from PBS injected mice, and significantly increased neutrophils present in the PEC of both zymosan-only and MSC-treated mice. However, examination of subsequent macrophage populations showed that whilst no macrophages were detected in the PBS control mice, both zymosan-only and CD317^pos^ MSC plus zymosan conditions displayed significant increases in macrophage numbers. Significantly fewer cells, including macrophages, were recruited in the presence of CD317^neg^ MSCs compared to zymosan only induction, therefore CD317^pos^ MSCs fail to inhibit macrophage recruitment.

The influence of CD317^pos^ MSCs on T cells appears to be highly modulated in comparison to CD317^neg^ MSCs. MSCs have been widely shown to deactivate T cells *in vitro* and suppress T cell proliferation whilst directing CD4+ effector T cells from Th1 to Th2 profile^66,78-85^. However, in activated T cells in cell to cell contact with CD317^pos^ MSCs, we observed minimal deactivation of T cells and continued T cell proliferation, in conjunction with an active increase in Th1 polarisation, contrary to the widely accepted immunosuppressive properties of MSCs. IFN-γ stimulation of MSCs has been shown to induce activation through upregulation of HLA class II, pushing the MSC towards antigen-presenting capability for immune regulation, promoting T cell interactions and potentially influencing CD8+ T cell activation^86^. This may go towards explaining the results we observe when CD317^pos^ cells interact with T cells *in vitro* and T and B cells *in vivo*. CD317^pos^ MSCs show minimal interaction with T cells *in vitro*, yet function more effectively in a pro-inflammatory *in vivo* environment. CD317 promotes an immune response through stimulating activation of NFκB^87^ which in turn contributes to B cell development^88^. MSC immunomodulation is intrinsically tied to interactions with dendritic cells (DCs), with MSCs inhibiting DC maturation, resulting in reduced migration, cytokine secretion, antigen presentation to T helper cells and cross-presentation to cytotoxic T cells^89^ through interrupting entry into the cell cycle, inhibiting DC differentiation and function^90^. DCs also mediate the MSC immunosuppressive effect through the induction of regulatory T cells^91,92^.

Deeper analysis of the CD317^pos^ subset of MSCs identified a heightened interferon signature that was not related to IFN-γ receptor expression levels, suggestive of constitutive IFN signalling. Pre-established, low level constitutive IFN signalling contributes to rapid pathogen responses in the innate immune system and conveys a protective effect to de novo IFN exposure in these cells^93^. CD317^pos^ MSCs, if maintained at appropriate levels, may therefore contribute to enhanced innate immunomodulation. Of interest, CD317^pos^ MSCs may also serve as a useful tool in the investigation of host tropism in viral infection, a particularly prevalent issue with the advent of COVID-19. Indeed, the presence of BST2/CD317 on the cell surface has been shown to convey a protective effect by tethering coronavirus virions to the cell surface or intracellular membranes and decreasing budding of progeny virus^94^. These cells may therefore provide an enhanced response to viral infection that facilitates tissue regeneration as well as immunomodulation. However, whilst constitutive IFN signalling may convey a protective effect to cells experiencing *de novo* IFN in the *in vivo* environment, there also exists the potential for a link between unregulated constitutive IFN signalling and tissue damage in human disease conditions including autoimmunity. It is therefore highly significant that we show the baseline gene expression levels of CD317^pos^ MSCs aligns them with cells present in autoimmune and related conditions.

In this report we characterise a subset of human MSCs that favour immunomodulatory interactions over tissue regeneration, yet identify as MSCs through both independent tests (e.g. Rohart) and ISCT guidelines^95^. These cells display a distinct immune profile and operate in contrast to the expectations of MSC’s immunosuppressive function. We have demonstrated that the proportion of CD317^pos^ MSCs varies considerably between donor MSC preparations, which could reflect individual inflammatory state and/or infection history. We propose that the success of therapeutic applications for tissue regeneration is dependent on the numbers of CD317^pos^ MSCs present in the administered cell dose. There is also the possibility that CD317^pos^ MSCs can bring therapeutic benefits in the inflamed environment. The expression of CD317 on MSCs serves as a positive marker for cells that display all the characteristics of an immune stromal cell and targeted therapies should aim to harness the knowledge of this cell type as novel approaches to the treatment of degenerative, and inflammatory conditions.

## Materials and Methods

### Cell culture

#### Immortalised MSC lines and primary bone marrow derived human MSCs

MSC lines immortalised with human telomerase reverse transcriptase (hTERT) were maintained in culture as previously described^60^. Clonal hTERT-MSCs included the CD317^pos^ Y202 and Y102 lines, and the CD317^neg^ Y201 and Y101 lines. Low-passage (p1-p5) primary MSCs were isolated from femoral heads, obtained with informed consent during routine hip replacement or as explant cultures from human tibial plateaux after routine knee replacement^60^. Primary MSCs were also established from bone marrow aspirates purchased from Lonza. Cells were cultured at 37°C in 5% CO_2_ humidified atmosphere incubaters using DMEM (Gibco) culture medium supplemented with 10% foetal bovine serum and 1% penicillin-streptomycin. Cells were routinely passaged at 80% confluence.

#### Isolation of primary T cells from tonsillectomy tissue

Primary donor T cells were retrieved from tonsillectomy donations according to ethical approval. For primary MSC co-cultures, cryopreserved CD4+ human cord blood T cells were purchased from Stem Cell Technologies. T cells were isolated from mixed T and B cell cultures using nylon wool separation^96^. T cells were seeded at a density of 1.0 × 10^6^ cells/ml in an appropriately sized tissue culture flask. MSC co-cultures with isolated T cells were set up within 24 hours or cells were cryopreserved in 10% dimethylsulfoxide (DMSO) in RPMI1640 medium and re-established in culture a minimum of 24 hours prior to use.

#### Rohart test for independent confirmation of MSC status

The Rohart MSC test was used as an independent measure for distinguishing MSCs from non-MSCs^61^. The classifier has previously been validated against 1,291 samples from 65 studies derived on 15 different platforms, with >95% accuracy with 97.7% accuracy^61^.

#### Flow cytometry

MSCs were labelled using optimised concentrations of the required primary antibody or isotype control. After washing, cells were stained with a fluorescent secondary antibody, where conjugated primaries were not used. As appropriate, cells were washed as required prior to incubation with 1:1000 diluted sytox blue for 5 minutes. Analysis was conducted immediately following staining.

Intracellular flow cytometry of MSC was performed on 4% paraformaldehyde (PFA) fixed cells in the presence of 0.1% saponin (Sigma). All flow cytometry was performed on a Beckman Coulter CyAn ADP flow cytometer and analysed with Summit v4.3 software, or using a Cytoflex S or LX and analysed with FCS Express 7. Cell sorting was undertaken using a Beckman Coulter MoFlo Astrios and analysed with summit v6.2 software or FCS Express 7.

#### Processing of mouse femurs

Femurs were dissected from C57BL/6J female mice at ages 8-12 weeks immediately after sacrificing. All work was carried out under ethical approval from the University of York Department of Biology Ethics Committee and Animal Welfare Ethical Review Body. Muscle tissue was removed and femurs were fixed in 4% PFA for 24 hours at 4°C, followed by washing with PBS. Bones were then decalcified using 10% EDTA in PBS at pH 7.5 for 24 hours at 4°C. After decalcification, femurs were cryoprotected by submerging in 30% sucrose in PBS for 24 hours at 4°C. Bones were embedded in Optimal Cutting Temperature compound and sectioned using an OTF5000 cryostat (Bright Instruments Ltd.). Sections were collected on SuperFrost plus microscope slides (Thermofisher) and stored at -70°C.

#### Immunofluorescent staining of mouse bone tissues

Slides were allowed to reach room temperature. Sections were blocked for 45 minutes in 10% goat serum (Sigma) + 0.1% Tween-20 in PBS (10% donkey serum (Sigma) + 0.1% Tween-20 in PBS where goat primary antibody was used). Primary antibodies (LEPR, CD31, CD317) were diluted in 1% IgG-free Bovine Serum Albumin (Sigma) + 0.05% Tween-20 (Sigma) in PBS and sections incubated in the dark at 4°C overnight in a humidified chamber. All secondary antibodies were added at 1:300 dilution in PBS for 1 hour at room temperature in the dark then stained for 10 minutes with 0.2 µg/ml 4′,6-diamidino-2-phenylindole (DAPI) in PBS. Dried slides were mounted with Prolong Gold antifade mounting medium (Invitrogen) and #1.5 thickness glass coverslip (Scientific Laboratory Supplies). Slides were left to cure at room temperature in the dark for 24 hours prior to image capture using LSM880 or LSM780 (Zeiss) confocal microscopes with excitation wavelengths of 405 nm, 488 nm, 561 nm and 633 nm.

#### Proteomic analysis of MSC plasma membranes

Plasma membranes were isolated from the hTERT immortalised clonal lines following the protocol of Holley *et al*^97^ before mass spectrometry and comparative proteomic analyses were performed by the Proteomics laboratory within the University of York Bioscience Technology Facility using LC-MS/MS^98^ and Scaffold 4 proteome software for initial analysis using 3% false discovery rate. Further in-depth examination of protein expression was conducted using the Knime analytics platform and ProteoWizard MSOpen technology^99^.

#### Transwell cell migration assays

Migration assays were performed in transwell polycarbonate membrane cell culture inserts with a 5µm pore (Corning, Sigma-Aldrich) using 1.25×10^5^ hTERT and primary MSCs, and monocyte-like THP-1 and T cell-like HUT-78 (ECACC 88041901) cells in 6 well plates with 1.5 ml of serum-free DMEM. After 24 hours, 600 µl of supernatant or DMEM was added in duplicate to the wells of the transwell plates. Polycarbonate filters were carefully placed above supernatant and 2.5×10^5^ of the appropriate cells in 100 µl serum-free RPMI-1640 were applied to the top of the filter and incubated for 5 hours before removing transwells. Migrated cells were assessed by flow cytometry. The percentage cells undergoing migration towards stimuli was calculated. For CCR2 testing, 500 nM CCR2 inhibitor was used (Teijin compound 1) in supernatant. Inhibition of migration was calculated as a percentage of cell total.

#### Examination of Gene Ontology (GO) terms in disease states for comparison with hTERT MSC lines

A bioinformatics comparison of the hTERT MSC lines gene expression data with publicly available transcriptomic data from a range of autoimmune and related disorders was undertaken to identify disease states that correlated with upregulated GO terms associated with the CD317^pos^ Y102 and Y202 clonal MSC lines^60^. Cross-platform validation was performed using Python and GeneSpring software was used to analyse outcomes. Differentially expressed genes were identified as greater than 2-fold upregulation in disease state compared to healthy controls, and GeneSpring was used to identify significance (p<0.05) in GO term occurrence. The 10 most upregulated GO terms were identified and comparisons made between autoimmune disease states and hTERT immortalised MSC lines.

#### Quantitative polymerase chain reaction (qPCR)

RNA was isolated from cells using TRIzol for cell lysis and Machery-Nagel RNA Nucleospin II kit for RNA isolation, with RNA converted to cDNA for gene expression analyses using Superscript IV reverse transcriptase enzymes (Invitrogen). Specific primers for gene expression analyses were designed and optimised and are described in Table S5. Gene expression analyses were performed as previously described^60^. Gene expression of eight IFN-γ regulated genes, namely *Ly6E, HERC5, IFI44L, ISG15, Mx1, Mx2, EPSTI1* and *RSAD2* were amplified in qPCR and fold changes were calculated relative to the expression of the housekeeping gene RPS27a and relative to the Y201 cell line or CD317^neg^ cells. The ΔΔCT fold changes were log2-transformed and averaged to calculate IFN-γ scores, as previously described^64,100^.

#### Enzyme-linked immunosorbent assays

To detect secreted proteins, supernatants from 100,000 cells incubated in 2.5 ml of serum free DMEM for 24 hours was analysed for secreted proteins by enzyme-linked immunosorbent assays (ELISA) using ELISA kits for CXCL10, CXCL11 (BioLegend); CCL2 (eBioscience); and SAA4 (Stratech) following manufacturers instructions.

#### PCR molecular diagnostics for infectious disease

Samples of hTERT lines Y201 and Y202 were tested externally and independently (Charles River) for viral contaminants using the Human Comprehensive cell line examination and report (CLEAR) Panel to detect RNA transcripts for 26 viral components, including virions commonly linked with autoimmune disorders (HIV, hepatitis, herpes simplex and herpesvirus, Epstein-Barr virus, BK virus, human T-Lymphotropic virus, Lymphocytic choriomeningitis virus and Cytomegalovirus)^101,102^. A low copy exogenous nucleic acid was added to sample lysis prior to nucleic acid isolation to serve as both a control to monitor for nucleic acid recovery and PCR inhibition. An RNA NRC was used to monitor reverse transcription for RNA virus assays. Nucleic acid recovery and PCR inhibition was monitored by a PCR assay specific for the NRC template.

#### T cell activation assay to assess MSC immunomodulation for deactivation and suppression of T cell proliferation

Co-culture of primary human tonsil T cells with hTERT MSC lines was used to assess the potential immunomodulatory impact of CD317^neg^ (Y101, Y201) and CD317^pos^ (Y102, Y202) cell lines on T cell proliferation and T helper differentiation. Continual proliferative capacity was used as a measure of T cell deactivation. hTERT MSC lines or CD317-sorted primary MSCs were seeded at a ratio of 1:10 with T cells with 1.0×10^4^ MSCs seeded into a 96-well U bottomed plate and cultured for 24 hours at 37°C, 5% CO_2_. Primary human MSC were sorted for CD317 expression and co-cultured with commercially sourced cryopreserved CD4+ human cord blood T cells (Stem Cell Technologies).

For assessment of proliferation, T cells were stained for 15 minutes at 37°C using 1 uM VPD450 Violet proliferation dye (eBioscience, Inc.). Unstained cells were used as a control. T cells were activated using anti-CD3ε/CD28 Dynabeads (Thermo Fisher) at a bead-to-cell ratio of 1:1 then seeded onto the MSC at a density of 1.0×10^5^/well (ratio 10:1) in 200 μl RPMI-1640 with 10% FBS, 0.05 μg/mL IL-2 (Peprotech, Inc) or seeded alone (no MSCs) as a control. Plates were cultured for 5 days at 37°C. T cell proliferation was assessed following removal of Dynabeads with the DynaMag-2 as per manufacturer’s recommendations. Plates were cultured for 5 days at 37°C. T cell proliferation was assessed with flow cytometry, with reduction in signal intensity visualised for repeated proliferation peaks. Proliferation was assessed through VPD450 dilution (diminished staining intensity) described through a proliferative index (PI) calculated from the fluorescence intensity at each cell division as described previously^70^. Proliferative cycles undertaken were calculated on 50% fluorescence intensity reduction peaks, measuring from fluorescence intensity of the first division and the final division detected.

#### T cell activation assay to assess MSC immunomodulation to direct effector T cell polarisation

For assessment of T helper differentiation, T cells were activated and cultured with hTERT MSC monolayers, as described above. The following reagents and antibodies for reactivation, transport inhibition and staining were sourced from eBioscience. Following 5 days of culture, T cells were re-stimulated using a combination of phorbol 12-myristate 13-acetate (PMA) (50 ng/ml) (Sigma Aldrich) and Ionomycin (1 μg/ml) (Invitrogen) and intracellular cytokines retained using transport inhibitor cocktail with 10 μg/ml brefeldin A and 2 μM Monensin (Invitrogen). Cells were cultured for 4 hours at 37°C then stained for surface marker CD4. Intracellular staining for helper T cells was undertaken for anti-human IFN-γ (Th1), IL-4 (Th2) or IL17a (Th17) or CD4 and CD25 then fixation/permeabilisation and staining for nuclear protein FOXP3 for regulatory T cells. All cells were measured using the CyAn ADP or Cytoflex LX flow cytometer and analysed with FCS Express 7. Comparisons were drawn for percentage of T helper differentiation within the CD4+ cell population and signal intensity (Median) for each antibody tested.

#### In vitro human skin explant model to assess cutaneous tissue damage

The human skin explant assay is an *in vitro* model previously used for evaluation of tissue damage induced by T cell or pro-inflammatory cytokine mediated immmunopathological responses^103,104^. We used this assay to investigate the *in situ* activities of CD317^neg^ Y201 and CD317^pos^ Y202 MSCs. Skin samples were obtained with informed consent and approval of the local research ethics committee (REC14/NE/1136, NRES Committee North East, IRAS project ID 129780). Following 48 hours stimulation with IFN-*γ* or TNF-*α* (both at 5 ng/ml), Y201 and Y202 MSCs were harvested, washed and plated at a density of 1×10^5^ cells/well in a 96 well round-bottomed plate. The cells were incubated for 3-4 hours to allow for adherence to the plastic. Two punch skin biopsies at 4 mm diameter taken from healthy volunteers were dissected into 10-12 sections of equal size. Each section was co-cultured with stimulated or unstimulated Y201 or Y202 in duplicate in a 200 μl total volume of DMEM supplemented with 20% heat–inactivated pooled human AB serum at 37°C and 5% CO_2_. Skin sections cultured in the culture medium containing 200 ng/ml IFN-γ or culture media alone were used as positive and background controls respectively. After 3-day culture, the skin sections were fixed in 10% formalin, then paraffin embedded and sectioned at 5 μm onto microscopic slides. The skin sections were stained with haematoxylin and eosin (H&E) following routine protocols. The severity of histopathological tissue damage was evaluated by two independent evaluators according to the Lerner scoring criteria^105^ as follows: grade 0, normal skin; grade I, mild vacuolization of epidermal basal cells; grade II, diffuse vacuolization of basal cells with scattered dyskeratotic bodies; grade III, subepidermal cleft formation; grade IV, complete epidermal separation^105^. Grade II or above were considered positive while Grade I changes considered as background, which is observed in skin sections cultured in medium alone.

#### In vivo assessment of immunomodulatory capacity of hTERT MSC lines in a murine peritonitis model

To determine the immunomodulatory properties of hTERT MSC lines, an *in vivo* zymosan-induced peritonitis model was used in C57BL/6J mice aged 8-10 weeks as described previously^106,107^. These experiments were carried out in accordance with the Animals and Scientific Procedures Act 1986, under UK Home Office Licence (project licence number PPL PFB579996 approved by the University of York Animal Welfare and Ethics Review Board). At day 0, mice were administered with an intraperitoneal infusion of 1 mg of zymosan A (Merck) in 100 μl of PBS. Immediately following the administration of zymosan, test condition mice were administered an intraperitoneal infusion of 2.0×10^6^ cells of either Y201 (CD317^neg^) or Y202 (CD317^pos^) in 100 μl of PBS; negative control mice were given PBS vehicle only.

After 24 hours, mice were euthanised using CO_2_ overdose and cervical dislocation. Intraperitoneal injection of 4 ml of ice cold RPMI-1640 was administered as peritoneal lavage. The process was repeated with a second 4 ml RPMI-1640 wash and wash solutions pooled to form the peritoneal exudate cells (PEC).

For each animal tested, red blood cells were lysed using Red Cell Lysis buffer (Merck) and a cell count performed. Spleens were retrieved from the mice and cell counts were recorded and a measure of spleen cellularity calculated. PEC samples were initially stained for Ly6C (APC), Ly6G (FITC), F4/80 (PE-Cy7) CD45 (PerCP-Cy5.5) (BioLegend) and Ly6G (FITC), CD11b (BUV395) and SiglecF (BV421) (BD). Both PEC and spleen samples were then stained for TCRb (AF488), CD3 (APC-Cy7), CD4 (PerCP-Cy5.5), CD62L (APC) and CD44 (PE) (BioLegend). Although at an early timepoint, spleen samples were additionally examined for T cell polarisation looking at T effector cells CD8 (PerCP-Cy5.5), CD4 (APC), IL4 (AF488), IFN-γ (PE) and IL17a (BV421) (BioLegend) and T reg cells using CD8 (PerCP-Cy5.5), CD4 (APC), CD25 (PE) and FOXP3 (AF488) (BioLegend). For all tests, Zombie Aqua (BioLegend) was used to exclude dead cells.

#### In vivo assay to assess tissue forming capacity of hTERT MSC lines

All procedures used were approved by the University of Leeds Ethics Committee and under the UK Home Office Project License (PPL:70/8549). The tissue-forming capacity of CD317^neg^ and CD317^pos^ hTERT cell lines CD317^neg^ Y201 and CD317^pos^ Y202 was assessed in CD1 nude mice (Charles River) aged 8-10 weeks in an *in vivo* transplantation assay^108^. 2.0 × 10^6^ MSC cell suspension in 1 ml medium was added to 40 mg hydroxyapatite (HA) synthetic bone particles (Zimmer Biomet) of 250-1000 μm size and rotated at approximately 25 rpm at 37°C for 100 minutes to allow cells to attach. HA particles were bound using fibrin glue comprising 30 μl thrombin (400 I.U./ml in DMEM medium) mixed 1:1 with fibrinogen (115 mg/ml in 0.85% saline solution). Implants were delivered subcutaneously into immunocompromised nude mice with two constructs placed into each mouse.

Transplants were harvested at 3 and 8 weeks, fixed in 4% PFA, decalcified for 7 days in 10% EDTA then stored overnight in 70% ethanol prior to paraffin embedding, sectioning and staining with H&E, Alcian Blue and Syrius Red (Thermo Fisher).

#### Statistical analysis

Data were tested for equal variance and normality using D’Agostino & Pearson omnibus normality test. Differences between groups were compared using two-tailed 1-way ANOVA for parametric data or Kruskall-Wallis for non-parametric testing. For two factor analysis, data was analysed with a two-tailed 2-way ANOVA. Bonferroni post-hoc testing was conducted to compare between groups. All statistical analysis was carried out using IBM SPSS Statistics 24.0, or GraphPad Prism version 5.0-9.0 with P<0.05 deemed statistically significant. Results are annotated as *p<0.05, **p<0.01, ***p<0.001 and all averaged values are expressed as mean ± standard error of the mean (SEM).

## Data Availability

Data will be made available in a publically accessible repository prior to publication.

## Author contributions

AGK designed, performed and analysed T cell experiments. AGK and JPH designed, performed and analysed peritonitis experiments. AS designed, performed and analysed MSC localisation experiments. JMF, SR and SJ designed, performed and analysed ELISA, Interferon signature, Rohart testing, cell migration experiments and bioinformatics. XY and EK performed subcutaneous HA scaffold implantation *in vivo* whilst AGK performed the associated cell culture and analysis of explants. PG designed experiments and was responsible for conceptualisation, funding acquisition, supervision and writing (review and editing). XW designed, performed and analysed the *in vitro* skin explant model. AK, JMF and PG wrote the paper.

## Acknowledgments

The authors thank the staff and patients of Clifton Park Hospital for samples. We are grateful to the University of York Technology Facility for support with flow cytometry, cell sorting, confocal microscopy bioinformatics and proteomic analysis. We thank Emily Taylor for assistance in isolating primary T cells. This work was funded by the Biotechnology and Biological Sciences Research Council (BBSRC) United Kingdom, Doctoral Training Partnership grant (BB/M011151/1) and the Tissue Engineering and Regenerative Therapies Centre Versus Arthritis (21156). LK and XY are partially funded by funding by the ‘EPSRC CDT in Tissue Engineering and Regenerative Medicine’ at the University of Leeds (Grant number EP/L014823/1). The York Centre of Excellence in Mass Spectrometry was created thanks to a major capital investment through Science City York, supported by Yorkshire Forward with funds from the Northern Way Initiative, and subsequent support from EPSRC (EP/K039660/1; EP/M028127/1).

## Materials & Correspondence

Requests and correspondence should be addressed to Professor Paul Genever.

## References

1 Pittenger, M. F. et al. Multilineage potential of adult human mesenchymal stem cells. Science 284, 143–147 (1999).

2 Colter, D. C., Class, R., DiGirolamo, C. M. & Prockop, D. J. Rapid expansion of recycling stem cells in cultures of plastic-adherent cells from human bone marrow. Proc Natl Acad Sci U S A 97, 3213–3218, doi:10.1073/pnas.070034097 (2000).

3 Bonab, M. M. et al. Aging of mesenchymal stem cell in vitro. BMC Cell Biol 7, 14, doi:10.1186/1471-2121-7-14 (2006).

4 Wilson, A., Hodgson-Garms, M., Frith, J. E. & Genever, P. Multiplicity of Mesenchymal Stromal Cells: Finding the Right Route to Therapy. Front Immunol 10, 1112, doi:10.3389/fimmu.2019.01112 (2019).

5 Najar, M. et al. Mesenchymal stromal cells and immunomodulation: A gathering of regulatory immune cells. Cytotherapy 18, 160–171, doi:10.1016/j.jcyt.2015.10.011 (2016).

6 Dominici, M. et al. Minimal criteria for defining multipotent mesenchymal stromal cells. The International Society for Cellular Therapy position statement. Cytotherapy 8, 315–317, doi:10.1080/14653240600855905 (2006).

7 Kuntin, D. & Genever, P. Mesenchymal stem cells from biology to therapy. Emerg Top Life Sci, doi:10.1042/ETLS20200303 (2021).

8 Wilson, A. J., Rand, E., Webster, A. J. & Genever, P. G. Characterisation of mesenchymal stromal cells in clinical trial reports: analysis of published descriptors. Stem Cell Research & Therapy 12, doi:10.1186/s13287-021-02435-1 (2021).

9 Prockop, D. J., Prockop, S. E. & Bertoncello, I. Are clinical trials with mesenchymal stem/progenitor cells too far ahead of the science? Lessons from experimental hematology. Stem cells 32, 3055–3061, doi:10.1002/stem.1806 (2014).

10 Wilson, A., Webster, A. & Genever, P. Nomenclature and heterogeneity: consequences for the use of mesenchymal stem cells in regenerative medicine. Regen Med 14, 595–611 (2019).

11 Plant, A. L. & Parker, G. C. Translating stem cell research from the bench to the clinic: a need for better quality data. Stem Cells Dev 22, 2457–2458, doi:10.1089/scd.2013.0188 (2013).

12 Jo, C. H. et al. Intra-articular injection of mesenchymal stem cells for the treatment of osteoarthritis of the knee: a proof-of-concept clinical trial. Stem Cells 32, 1254–1266, doi:10.1002/stem.1634 (2014).

13 Migliorini, F. et al. Improved outcomes after mesenchymal stem cells injections for knee osteoarthritis: results at 12-months follow-up: a systematic review of the literature. Arch Orthop Trauma Surg 140, 853–868, doi:10.1007/s00402-019-03267-8 (2020).

14 Ma, W. et al. Efficacy and safety of intra-articular injection of mesenchymal stem cells in the treatment of knee osteoarthritis: A systematic review and meta-analysis. Medicine (Baltimore) 99, e23343, doi:10.1097/MD.0000000000023343 (2020).

15 Gupta, P. K. et al. Efficacy and safety of adult human bone marrow-derived, cultured, pooled, allogeneic mesenchymal stromal cells (Stempeucel(R)): preclinical and clinical trial in osteoarthritis of the knee joint. Arthritis Res Ther 18, 301, doi:10.1186/s13075-016-1195-7 (2016).

16 Shim, J. et al. Safety and efficacy of Wharton’s jelly-derived mesenchymal stem cells with teriparatide for osteoporotic vertebral fractures: A phase I/IIa study. Stem Cells Transl Med 10, 554–567, doi:10.1002/sctm.20-0308 (2021).

17 Wang, L. et al. Efficacy and Safety of Umbilical Cord Mesenchymal Stem Cell Therapy for Rheumatoid Arthritis Patients: A Prospective Phase I/II Study. Drug Des Devel Ther 13, 4331–4340, doi:10.2147/DDDT.S225613 (2019).

18 El-Jawhari, J. J., El-Sherbiny, Y., McGonagle, D. & Jones, E. Multipotent Mesenchymal Stromal Cells in Rheumatoid Arthritis and Systemic Lupus Erythematosus; From a Leading Role in Pathogenesis to Potential Therapeutic Saviors? Front Immunol 12, 643170, doi:10.3389/fimmu.2021.643170 (2021).

19 Alvaro-Gracia, J. M. et al. Intravenous administration of expanded allogeneic adipose-derived mesenchymal stem cells in refractory rheumatoid arthritis (Cx611): results of a multicentre, dose escalation, randomised, single-blind, placebo-controlled phase Ib/IIa clinical trial. Ann Rheum Dis 76, 196–202, doi:10.1136/annrheumdis-2015-208918 (2017).

20 Carlsson, P. O., Schwarcz, E., Korsgren, O. & Le Blanc, K. Preserved beta-cell function in type 1 diabetes by mesenchymal stromal cells. Diabetes 64, 587–592, doi:10.2337/db14-0656 (2015).

21 Lin, W. et al. Administration of mesenchymal stem cells in diabetic kidney disease: a systematic review and meta-analysis. Stem Cell Res Ther 12, 43, doi:10.1186/s13287-020-02108-5 (2021).

22 Karussis, D. et al. Safety and immunological effects of mesenchymal stem cell transplantation in patients with multiple sclerosis and amyotrophic lateral sclerosis. Arch Neurol 67, 1187–1194, doi:10.1001/archneurol.2010.248 (2010).

23 Petrou, P. et al. Beneficial effects of autologous mesenchymal stem cell transplantation in active progressive multiple sclerosis. Brain 143, 3574–3588, doi:10.1093/brain/awaa333 (2020).

24 Lin, B. L. et al. Allogeneic bone marrow-derived mesenchymal stromal cells for hepatitis B virus-related acute-on-chronic liver failure: A randomized controlled trial. Hepatology 66, 209–219, doi:10.1002/hep.29189 (2017).

25 Casiraghi, F. et al. Third-party bone marrow-derived mesenchymal stromal cell infusion before liver transplantation: A randomized controlled trial. Am J Transplant, doi:10.1111/ajt.16468 (2020).

26 Suk, K. T. et al. Transplantation with autologous bone marrow-derived mesenchymal stem cells for alcoholic cirrhosis: Phase 2 trial. Hepatology 64, 2185–2197, doi:10.1002/hep.28693 (2016).

27 Mazzini, L. et al. Mesenchymal stromal cell transplantation in amyotrophic lateral sclerosis: a long-term safety study. Cytotherapy 14, 56–60, doi:10.3109/14653249.2011.613929 (2012).

28 Berry, J. D. et al. NurOwn, phase 2, randomized, clinical trial in patients with ALS: Safety, clinical, and biomarker results. Neurology 93, e2294–e2305, doi:10.1212/WNL.0000000000008620 (2019).

29 Staff, N. P. et al. Safety of intrathecal autologous adipose-derived mesenchymal stromal cells in patients with ALS. Neurology 87, 2230–2234, doi:10.1212/WNL.0000000000003359 (2016).

30 Oh, K. W. et al. Repeated Intrathecal Mesenchymal Stem Cells for Amyotrophic Lateral Sclerosis. Ann Neurol 84, 361–373, doi:10.1002/ana.25302 (2018).

31 Shi, L. et al. Effect of human umbilical cord-derived mesenchymal stem cells on lung damage in severe COVID-19 patients: a randomized, double-blind, placebo-controlled phase 2 trial. Signal Transduct Target Ther 6, 58, doi:10.1038/s41392-021-00488-5 (2021).

32 Chouw, A. et al. Potency of Mesenchymal Stem Cell and Its Secretome in Treating COVID-19. Regen Eng Transl Med, 1–12, doi:10.1007/s40883-021-00202-5 (2021).

33 Leng, Z. et al. Transplantation of ACE2(-) Mesenchymal Stem Cells Improves the Outcome of Patients with COVID-19 Pneumonia. Aging Dis 11, 216–228, doi:10.14336/AD.2020.0228 (2020).

34 Lalu, M. M. et al. Safety of cell therapy with mesenchymal stromal cells (SafeCell): a systematic review and meta–analysis of clinical trials. PLoS One 7, e47559, doi:10.1371/journal.pone.0047559 (2012).

35 Majore, I., Moretti, P., Hass, R. & Kasper, C. Identification of subpopulations in mesenchymal stem cell-like cultures from human umbilical cord. Cell Commun Signal 7, 6, doi:10.1186/1478-811X-7-6 (2009).

36 Gullo, F. & De Bari, C. Prospective purification of a subpopulation of human synovial mesenchymal stem cells with enhanced chondro-osteogenic potency. Rheumatology (Oxford) 52, 1758–1768, doi:10.1093/rheumatology/ket205 (2013).

37 Yang, Z. X. et al. CD106 identifies a subpopulation of mesenchymal stem cells with unique immunomodulatory properties. PLoS One 8, e59354, doi:10.1371/journal.pone.0059354 (2013).

38 Blazquez-Martinez, A. et al. c-Kit identifies a subpopulation of mesenchymal stem cells in adipose tissue with higher telomerase expression and differentiation potential. Differentiation 87, 147–160, doi:10.1016/j.diff.2014.02.007 (2014).

39 Mo, M., Wang, S., Zhou, Y., Li, H. & Wu, Y. Mesenchymal stem cell subpopulations: phenotype, property and therapeutic potential. Cell Mol Life Sci 73, 3311–3321, doi:10.1007/s00018-016-2229-7 (2016).

40 Li, N. et al. The Sca1+ mesenchymal stromal subpopulation promotesdendritic cell commitment in the niche. Turkish Journal of Biology 41, 58–65, doi:10.3906/biy-1510-81 (2017).

41 Friedenstein, A. J. et al. Precursors for fibroblasts in different populations of hematopoietic cells as detected by the in vitro colony assay method. Exp Hematol 2, 83–92 (1974).

42 Friedenstein, A. J., Gorskaja, J. F. & Kulagina, N. N. Fibroblast precursors in normal and irradiated mouse hematopoietic organs. Exp Hematol 4, 267–274 (1976).

43 Tondreau, T. et al. Mesenchymal stem cells derived from CD133-positive cells in mobilized peripheral blood and cord blood: proliferation, Oct4 expression, and plasticity. Stem Cells 23, 1105–1112, doi:10.1634/stemcells.2004-0330 (2005).

44 Eirin, A. et al. Adipose tissue-derived mesenchymal stem cells improve revascularization outcomes to restore renal function in swine atherosclerotic renal artery stenosis. Stem Cells 30, 1030–1041, doi:10.1002/stem.1047 (2012).

45 Zuk, P. A. et al. Human adipose tissue is a source of multipotent stem cells. Mol Biol Cell 13, 4279–4295, doi:10.1091/mbc.e02-02-0105 (2002).

46 De Bari, C., Dell’Accio, F., Tylzanowski, P. & Luyten, F. P. Multipotent mesenchymal stem cells from adult human synovial membrane. Arthritis Rheum 44, 1928–1942, doi:10.1002/1529-0131(200108)44:8<1928::AID-ART331>3.0.CO;2-P (2001).

47 Hermida-Gomez, T. et al. Quantification of cells expressing mesenchymal stem cell markers in healthy and osteoarthritic synovial membranes. J Rheumatol 38, 339–349, doi:10.3899/jrheum.100614 (2011).

48 Gronthos, S., Mankani, M., Brahim, J., Robey, P. G. & Shi, S. Postnatal human dental pulp stem cells (DPSCs) in vitro and in vivo. Proc Natl Acad Sci U S A 97, 13625–13630, doi:10.1073/pnas.240309797 (2000).

49 Schwab, K. E., Hutchinson, P. & Gargett, C. E. Identification of surface markers for prospective isolation of human endometrial stromal colony-forming cells. Hum Reprod 23, 934–943, doi:10.1093/humrep/den051 (2008).

50 Park, J. C. et al. Isolation and characterization of human periodontal ligament (PDL) stem cells (PDLSCs) from the inflamed PDL tissue: in vitro and in vivo evaluations. J Clin Periodontol 38, 721–731, doi:10.1111/j.1600-051X.2011.01716.x (2011).

51 Bi, Y. et al. Identification of tendon stem/progenitor cells and the role of the extracellular matrix in their niche. Nat Med 13, 1219–1227, doi:10.1038/nm1630 (2007).

52 Noth, U. et al. Multilineage mesenchymal differentiation potential of human trabecular bone-derived cells. J Orthop Res 20, 1060–1069, doi:10.1016/S0736-0266(02)00018-9 (2002).

53 Romanov, Y. A., Svintsitskaya, V. A. & Smirnov, V. N. Searching for alternative sources of postnatal human mesenchymal stem cells: candidate MSC-like cells from umbilical cord. Stem Cells 21, 105–110, doi:10.1634/stemcells.21-1-105 (2003).

54 Baksh, D., Yao, R. & Tuan, R. S. Comparison of proliferative and multilineage differentiation potential of human mesenchymal stem cells derived from umbilical cord and bone marrow. Stem Cells 25, 1384–1392, doi:10.1634/stemcells.2006-0709 (2007).

55 Sarugaser, R., Lickorish, D., Baksh, D., Hosseini, M. M. & Davies, J. E. Human umbilical cord perivascular (HUCPV) cells: a source of mesenchymal progenitors. Stem Cells 23, 220–229, doi:10.1634/stemcells.2004-0166 (2005).

56 Martin-Rendon, E. et al. 5-Azacytidine-treated human mesenchymal stem/progenitor cells derived from umbilical cord, cord blood and bone marrow do not generate cardiomyocytes in vitro at high frequencies. Vox Sang 95, 137–148, doi:10.1111/j.1423-0410.2008.01076.x (2008).

57 Fukuchi, Y. et al. Human placenta-derived cells have mesenchymal stem/progenitor cell potential. Stem Cells 22, 649–658, doi:10.1634/stemcells.22-5-649 (2004).

58 Meirelles, L. D. S., Chagastelles, P. C. & Nardi, N. B. Mesenchymal stem cells reside in virtually all post-natal organs and tissues. Journal of Cell Science 119, 2204–2213, doi:10.1242/jcs.02932 (2006).

59 Crisan, M. et al. A perivascular origin for mesenchymal stem cells in multiple human organs. Cell Stem Cell 3, 301–313, doi:10.1016/j.stem.2008.07.003 (2008).

60 James, S. et al. Multiparameter Analysis of Human Bone Marrow Stromal Cells Identifies Distinct Immunomodulatory and Differentiation-Competent Subtypes. Stem Cell Reports 4, 1004–1015, doi:10.1016/j.stemcr.2015.05.005 (2015).

61 Rohart, F. et al. A molecular classification of human mesenchymal stromal cells. PeerJ 4, e1845, doi:10.7717/peerj.1845 (2016).

62 Balzano, M. et al. Nidogen-1 Contributes to the Interaction Network Involved in Pro-B Cell Retention in the Peri-sinusoidal Hematopoietic Stem Cell Niche. Cell Rep 26, 3257–3271 e3258, doi:10.1016/j.celrep.2019.02.065 (2019).

63 Zhou, B. O., Yue, R., Murphy, M. M., Peyer, J. G. & Morrison, S. J. Leptin-receptor-expressing mesenchymal stromal cells represent the main source of bone formed by adult bone marrow. Cell Stem Cell 15, 154–168, doi:10.1016/j.stem.2014.06.008 (2014).

64 Raterman, H. G. et al. The interferon type I signature towards prediction of non-response to rituximab in rheumatoid arthritis patients. Arthritis research & therapy 14, R95, doi:10.1186/ar3819 (2012).

65 Mostafavi, S. et al. Parsing the Interferon Transcriptional Network and Its Disease Associations. Cell 164, 564–578, doi:10.1016/j.cell.2015.12.032 (2016).

66 Kay, A. G. et al. Mesenchymal Stem Cell-Conditioned Medium Reduces Disease Severity and Immune Responses in Inflammatory Arthritis. Sci Rep 7, 18019, doi:10.1038/s41598-017-18144-w (2017).

67 Ren, G. et al. Mesenchymal stem cell-mediated immunosuppression occurs via concerted action of chemokines and nitric oxide. Cell Stem Cell 2, 141–150, doi:10.1016/j.stem.2007.11.014 (2008).

68 Keating, A. Mesenchymal stromal cells. Current Opinion in Hematology 13, 419–425, doi:10.1097/01.moh.0000245697.54887.6f (2006).

69 Nauta, A. J. & Fibbe, W. E. Immunomodulatory properties of mesenchymal stromal cells. Blood 110, 3499–3506, doi:10.1182/blood-2007-02-069716 (2007).

70 Angulo, R. & Fulcher, D. A. Measurement of Candida-specific blastogenesis: comparison of carboxyfluorescein succinimidyl ester labelling of T cells, thymidine incorporation, and CD69 expression. Cytometry 34, 143–151 (1998).

71 Que, J., Lian, Q., El Oakley, R. M., Lim, B. & Lim, S. K. PI3 K/Akt/mTOR-mediated translational control regulates proliferation and differentiation of lineage-restricted RoSH stem cell lines. J Mol Signal 2, 9, doi:10.1186/1750-2187-2-9 (2007).

72 Sivanathan, K. N. et al. Interleukin-17A-Induced Human Mesenchymal Stem Cells Are Superior Modulators of Immunological Function. Stem Cells 33, 2850–2863, doi:10.1002/stem.2075 (2015).

73 Bibby, L., Ribeiro, A., Ahmad, S. F. & Dickinson, A. M. A novel in-vitro human skin explant test to predict adverse immune reactions to biologics and aggregated monoclonal antibodies. Toxicology Letters 295, S66–S67, doi:10.1016/j.toxlet.2018.06.072 (2018).

74 Dickinson, A., Wang, X. N. & Ahmed, S. in Alternatives for Dermal Toxicity Testing (eds Chantra Eskes, Erwin van Vliet, & Howard I. Maibach) 437–448 (Springer International Publishing, 2017).

75 Wynn, T. A. & Vannella, K. M. Macrophages in Tissue Repair, Regeneration, and Fibrosis. Immunity 44, 450–462, doi:10.1016/j.immuni.2016.02.015 (2016).

76 Shechter, R. et al. Recruitment of beneficial M2 macrophages to injured spinal cord is orchestrated by remote brain choroid plexus. Immunity 38, 555–569, doi:10.1016/j.immuni.2013.02.012 (2013).

77 Butterfield, T. A., Best, T. M. & Merrick, M. A. The Dual Roles of Neutrophils and Macrophages in Inflammation: A Critical Balance Between Tissue Damage and Repair. Journal of Athletic Training 41, 457–465 (2006).

78 Augello, A. et al. Bone marrow mesenchymal progenitor cells inhibit lymphocyte proliferation by activation of the programmed death 1 pathway. Eur J Immunol 35, 1482–1490, doi:10.1002/eji.200425405 (2005).

79 Bloom, D. D. et al. A reproducible immunopotency assay to measure mesenchymal stromal cell-mediated T-cell suppression. Cytotherapy 17, 140–151, doi:10.1016/j.jcyt.2014.10.002 (2015).

80 Chinnadurai, R., Copland, I. B., Patel, S. R. & Galipeau, J. IDO-independent suppression of T cell effector function by IFN-gamma-licensed human mesenchymal stromal cells. J Immunol 192, 1491–1501, doi:10.4049/jimmunol.1301828 (2014).

81 Cuerquis, J. et al. Human mesenchymal stromal cells transiently increase cytokine production by activated T cells before suppressing T-cell proliferation: effect of interferon-gamma and tumor necrosis factor-alpha stimulation. Cytotherapy 16, 191–202, doi:10.1016/j.jcyt.2013.11.008 (2014).

82 Di Nicola, M. et al. Human bone marrow stromal cells suppress T-lymphocyte proliferation induced by cellular or nonspecific mitogenic stimuli. Blood 99, 3838–3843, doi:10.1182/blood.v99.10.3838 (2002).

83 Saldanha-Araujo, F. et al. Mesenchymal stromal cells up-regulate CD39 and increase adenosine production to suppress activated T-lymphocytes. Stem Cell Res 7, 66–74, doi:10.1016/j.scr.2011.04.001 (2011).

84 Sattler, C. et al. Inhibition of T-cell proliferation by murine multipotent mesenchymal stromal cells is mediated by CD39 expression and adenosine generation. Cell Transplant 20, 1221–1230, doi:10.3727/096368910X546553 (2011).

85 Tobin, L. M., Healy, M. E., English, K. & Mahon, B. P. Human mesenchymal stem cells suppress donor CD4(+) T cell proliferation and reduce pathology in a humanized mouse model of acute graft-versus-host disease. Clin Exp Immunol 172, 333–348, doi:10.1111/cei.12056 (2013).

86 van Megen, K. M. et al. Activated Mesenchymal Stromal Cells Process and Present Antigens Regulating Adaptive Immunity. Front Immunol 10, 694, doi:10.3389/fimmu.2019.00694 (2019).

87 Hotter, D., Sauter, D. & Kirchhoff, F. Emerging role of the host restriction factor tetherin in viral immune sensing. J Mol Biol 425, 4956–4964, doi:10.1016/j.jmb.2013.09.029 (2013).

88 Ishikawa, J. et al. Molecular Cloning and Chromosomal Mapping of a Bone Marrow Stromal Cell Surface Gene, BSTZ, That May Be Involved in Pre-B-Cell Growth. Genomics 26, 527–534 (1995).

89 Chiesa, S. et al. Mesenchymal stem cells impair in vivo T-cell priming by dendritic cells. Proc Natl Acad Sci U S A 108, 17384–17389, doi:10.1073/pnas.1103650108 (2011).

90 Ramasamy, R. et al. Mesenchymal stem cells inhibit dendritic cell differentiation and function by preventing entry into the cell cycle. Transplantation 83, 71–76, doi:10.1097/01.tp.0000244572.24780.54 (2007).

91 Choi, Y. S., Jeong, J. A. & Lim, D. S. Mesenchymal stem cell-mediated immature dendritic cells induce regulatory T cell-based immunosuppressive effect. Immunol Invest 41, 214–229, doi:10.3109/08820139.2011.619022 (2012).

92 Zhao, Z. G. et al. Immunomodulatory function of regulatory dendritic cells induced by mesenchymal stem cells. Immunol Invest 41, 183–198, doi:10.3109/08820139.2011.607877 (2012).

93 Sarhan, J. et al. Constitutive interferon signaling maintains critical threshold of MLKL expression to license necroptosis. Cell Death Differ 26, 332–347, doi:10.1038/s41418-018-0122-7 (2019).

94 Wang, S. M., Huang, K. J. & Wang, C. T. BST2/CD317 counteracts human coronavirus 229E productive infection by tethering virions at the cell surface. Virology 449, 287–296, doi:10.1016/j.virol.2013.11.030 (2014).

95 Friedenstein, A. J., Piatetzky-Shapiro, I. & Petrakova, K. V. Osteogenesis in transplants of bone marrow. Journal of Embryology and Experimental Morphology 16, 581–390 (1966).

96 Trizio, D. & Cudkowicz, G. Separation of T and B Lymphocytes by Nylon Wool Columns: Evaluation of Efficacy by Functional Assays in Vivo. The Journal of Immunology 113, 1093–1097 (1974).

97 Holley, R. J. et al. Comparative quantification of the surfaceome of human multipotent mesenchymal progenitor cells. Stem cell reports 4, 473–488, doi:10.1016/j.stemcr.2015.01.007 (2015).

98 Dowle, A. A., Wilson, J. & Thomas, J. R. Comparing the Diagnostic Classification Accuracy of iTRAQ, Peak-Area, Spectral-Counting, and emPAI Methods for Relative Quantification in Expression Proteomics. J Proteome Res 15, 3550–3562, doi:10.1021/acs.jproteome.6b00308 (2016).

99 Berthold, M. R. et al. Knime: The Konstanz Information Miner. (Springer, 2007).

100 de Jong, T. D. et al. Effect of prednisone on type I interferon signature in rheumatoid arthritis: consequences for response prediction to rituximab. Arthritis research & therapy 17, 78, doi:10.1186/s13075-015-0564-y (2015).

101 Smatti, M. K. et al. Viruses and Autoimmunity: A Review on the Potential Interaction and Molecular Mechanisms. Viruses 11, doi:10.3390/v11080762 (2019).

102 Sundsfjord, A. et al. BK and JC viruses in patients with systemic lupus erythematosus: prevalent and persistent BK viruria, sequence stability of the viral regulatory regions, and nondetectable viremia. J Infect Dis 180, 1–9, doi:10.1086/314830 (1999).

103 Dickinson, A. M., Sviland, L., Dunn, J., Carey, P. & Proctor, S. J. Demonstration of direct involvement of cytokines in graft-versus-host reactions using an in vitro human skin explant model. Bone Marrow Transplant 7, 209–216 (1991).

104 Dickinson, A. M. et al. In situ dissection of the graft-versus-host activities of cytotoxic T cells specific for minor histocompatibility antigens. Nat Med 8, 410–414, doi:10.1038/nm0402-410 (2002).

105 Lerner, K. G. et al. Histopathology of graft-vs.-host reaction (GvHR) in human recipients of marrow from HL-A-matched sibling donors. Transplant Proc 6, 367–371 (1974).

106 Choi, H., Lee, R. H., Bazhanov, N., Oh, J. Y. & Prockop, D. J. Anti-inflammatory protein TSG-6 secreted by activated MSCs attenuates zymosan-induced mouse peritonitis by decreasing TLR2/NF-kappaB signaling in resident macrophages. Blood 118, 330–338, doi:10.1182/blood-2010-12-327353 (2011).

107 Bianco, P., Kuznetsov, S. A., Riminucci, M. & Gehron Robey, P. in Adult Stem Cells Methods in Enzymology 117–148 (2006).

108 Kuznetsov, S. A. et al. The interplay of osteogenesis and hematopoiesis: expression of a constitutively active PTH/PTHrP receptor in osteogenic cells perturbs the establishment of hematopoiesis in bone and of skeletal stem cells in the bone marrow. J Cell Biol 167, 1113–1122, doi:10.1083/jcb.200408079 (2004).

