## Supplementary Figures and Tables for "Identification of CD317-Positive Pro-inflammatory Immune Stromal Cells in Human Mesenchymal Stromal Cell Preparations"

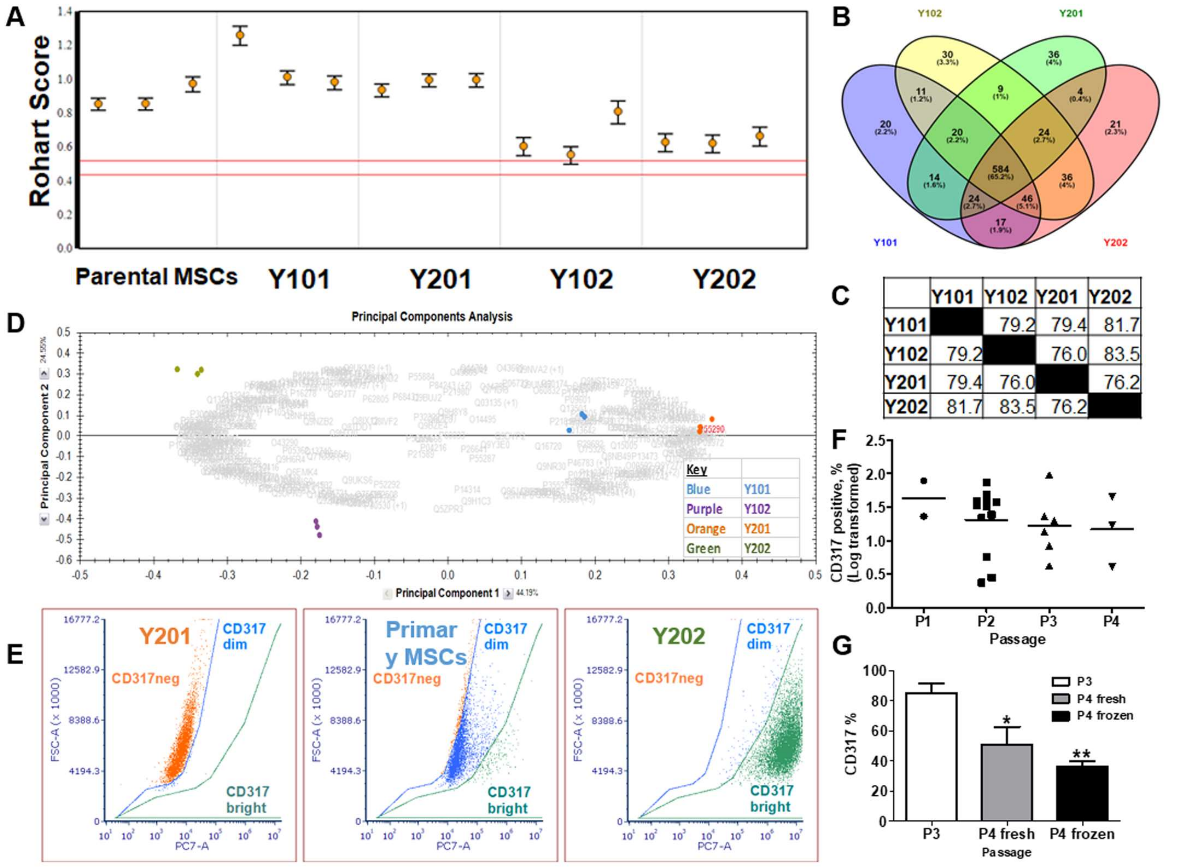


**Figure S1.** (A) Independent testing confirms MSC status for all immortalised MSC lines (Y101, Y201, Y102, Y202) using the Rohart Test. (B) Proteomic analysis of membrane proteins in MSC lines identified a small number of unique membrane proteins and (C) a high percentage of protein expression similarity between lines showing similar stromal proteomic profiles. (D) Principle component analysis (PCA) of mass spectrometry data identified distinct MSC populations, with Y102 and Y202 lines lying further from the population mean. (E) Representative flow cytometric analysis of primary MSCs shows the presence of CD317$^{neg}$ and CD317$^{pos}$ cells where CD317 positivity can be delineated into CD317$^{dim}$ and CD317$^{bright}$ (Y102 and Y202 equivalent). (F) CD317 expression over time in culture (passages 1-4, n=2, 12, 7, 3) is variable due to initial proportions of CD317$^{pos}$ cells but log transformed data shows no significant change over time ($p>0.05$, 1 Way ANOVA with Bonferroni post hoc test). (G) The effect of freeze cycles on cells demonstrates a significant reduction in CD317 expression due to cell freezing (n=5, 1 Way ANOVA with Bonferroni post hoc test). *$p<0.05$, **$p<0.01$.

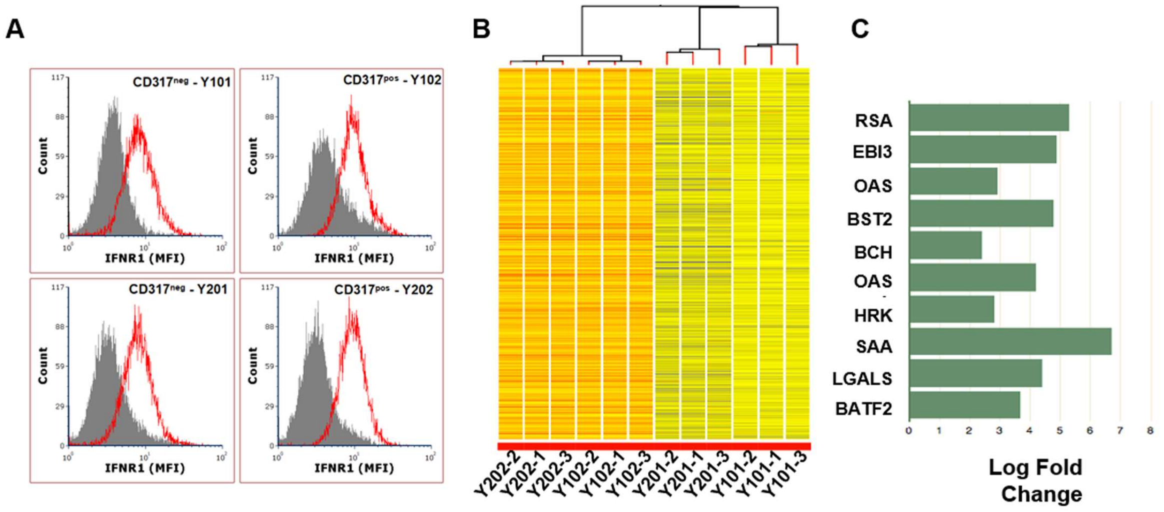


**Figure S2.** A) Representative histograms (of staining performed on two passages of MSC lines) of IFN-γ receptor type I cell surface expression by flow cytometry. B) Differentially expressed genes (DEGs) between Y102/Y202 and Y101/Y201 samples, hierarchical clustering of 2340 significantly upregulated DEGs in Y02 MSC samples (FC>2, $p<0.05$) with clustering of the Y01 group (Y101, Y201) and the Y02 group (Y102, Y202). C) The ten most significantly upregulated genes in the Y02 group versus Y01 group ($p<0.05$).

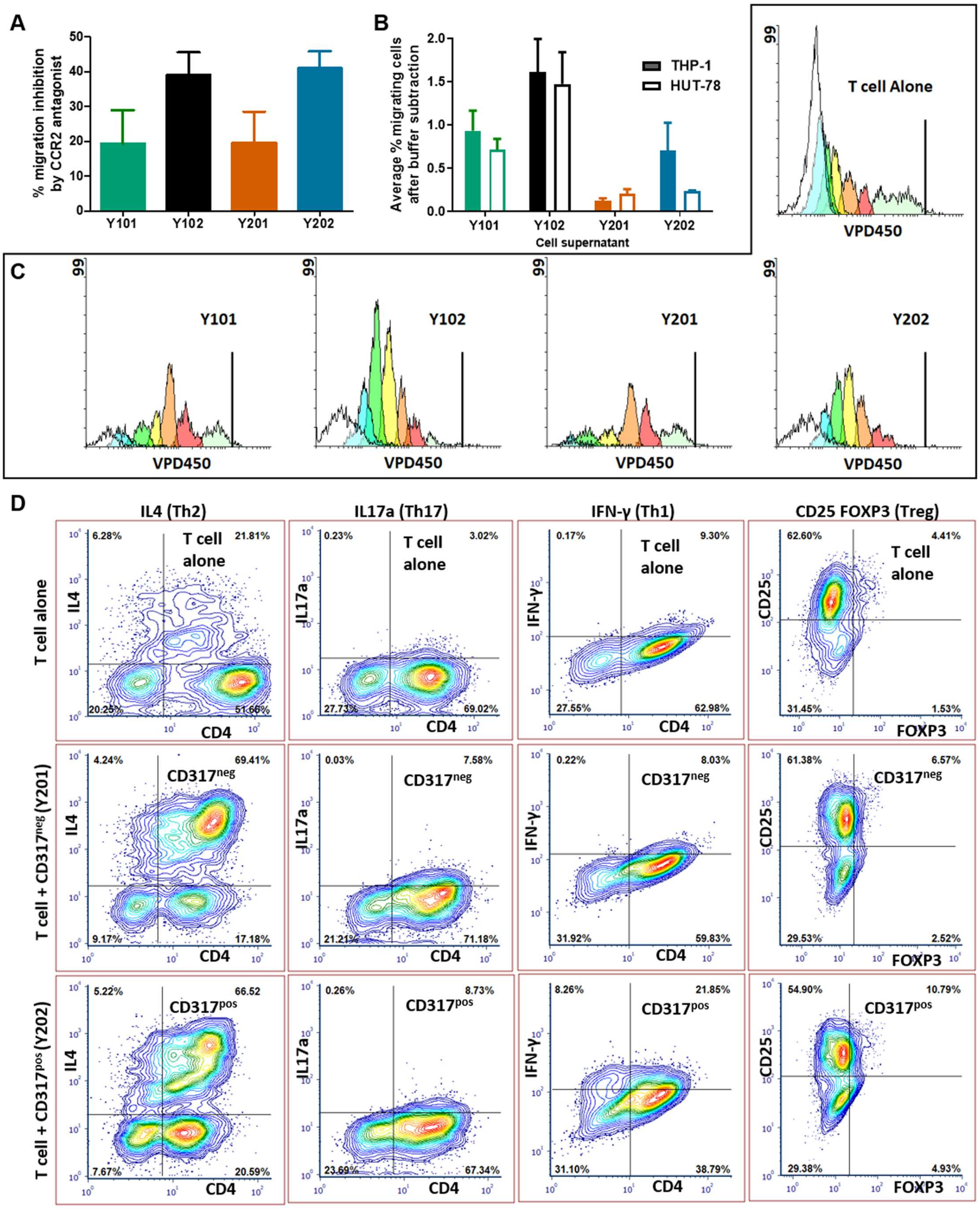


**Figure S3.** (A) Mean ± SEM percentage inhibition of monocyte migration induced by cell line supernatant of a CCR2 antagonist (n=4; experiments performed in duplicate). (B) Percentage inhibition of monocyte migration induced by cell line supernatant of a CCR2 antagonist. (C) Representative proliferation cycles as demonstrated by flow cytometry for hTERT immortalised CD317$^{neg}$ (Y101, Y201) and CD317$^{pos}$ (Y102, Y202) MSC lines. (D) Representative flow cytometry dot plots showing differentiation of T cells following polyclonal activation and culture alone, with Y201 cells or with Y202 cells.

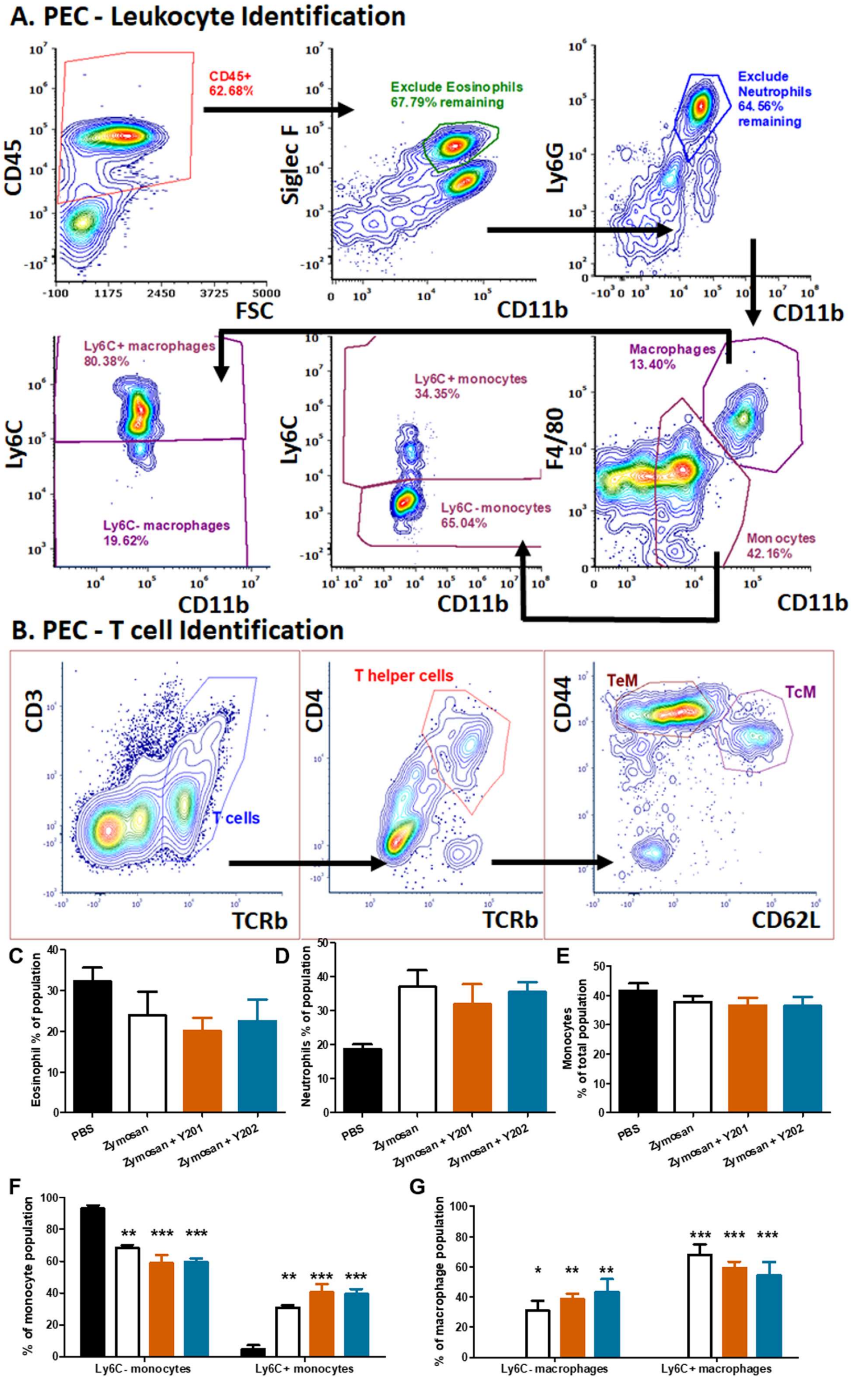


**Figure S4.** (A, B) Gating strategy was devised for flow cytometric analysis of multiple PEC cell types focused on haematopoietic, myeloid and lymphoid cells including monocytes, macrophages and T cells. (C, D, E) No difference was observed in recruitment of eosinophils, neutrophils or monocytes in MSC-treated mice when compared to zymosan alone or PBS controls. (F, G) Within monocyte and macrophage populations, proportions of Ly6C positive and negative cells matched the proportions seen in zymosan treatment only animals.

**Supplementary Tables**

Index:

Table S1 – Primer Sequences

Table S2 – Rohart Scores for individual cell lines

Table S3 – Gene Expression Omnibus (GEO) summary data for 01 and 02 cell lines

Table S4 – Gene ontology (GO) term associations between CD317$^{pos}$ cell lines and autoimmune disease states

Table S5 – Upregulated signalling pathway associations between CD317$^{pos}$ cell lines and autoimmune disease states

**Table S1. Rohart scores for parental cell line FH181 and hTERTimmortalised cells**

| Replicate Group ID | Prediction | Lower Bound | Upper Bound | MSC Calls | Total Sub-samplings | Percentage |
|---|---|---|---|---|---|---|
| FH181, replicate 1 | 0.85 | 0.82 | 0.89 | 200 | 200 | 100 |
| FH181, replicate 2 | 0.86 | 0.82 | 0.89 | 200 | 200 | 100 |
| FH181, replicate 3 | 0.97 | 0.92 | 1.01 | 200 | 200 | 100 |
| Y101, replicate 1 | 1.26 | 1.2 | 1.31 | 200 | 200 | 100 |
| Y101, replicate 2 | 1.01 | 0.97 | 1.05 | 200 | 200 | 100 |
| Y101, replicate 3 | 0.98 | 0.94 | 1.02 | 200 | 200 | 100 |
| Y102, replicate 1 | 0.6 | 0.55 | 0.65 | 200 | 200 | 100 |
| Y102, replicate 2 | 0.55 | 0.5 | 0.6 | 179 | 200 | 89.5 |
| Y102, replicate 3 | 0.81 | 0.74 | 0.87 | 200 | 200 | 100 |
| Y201, replicate 1 | 0.94 | 0.9 | 0.97 | 200 | 200 | 100 |
| Y201, replicate 2 | 1 | 0.95 | 1.03 | 200 | 200 | 100 |
| Y201, replicate 3 | 1 | 0.95 | 1.03 | 200 | 200 | 100 |
| Y202, replicate 1 | 0.63 | 0.57 | 0.68 | 200 | 200 | 100 |
| Y202, replicate 2 | 0.62 | 0.57 | 0.67 | 200 | 200 | 100 |
| Y202, replicate 3 | 0.66 | 0.6 | 0.72 | 200 | 200 | 100 |

**Table S2. Gene Expression Omnibus (GEO) data summary.**

Six datasets including Y02 group data were used. 'Sample 1' column states the disease condition and the 'Sample 2' column shows their relevant controls to which their upregulated gene expression was compared. Cell types analyzed, number of samples in each dataset, and the platform type they were analyzed on are shown.

| Sample 1 (S1) | Sample 2 (S2) | Cell Type | Sample ratio (S1:S2) | Platform Type | GEO Accession |
|---|---|---|---|---|---|
| **Y02** | **Y01** | Bone marrow MSCs | 6:6 | GPL14550 Agilent-028004 | - |
| **Psoriasis** | **Healthy Control** | Skin tissue | 25:9 | GPL14550 Agilent-028004 | GSE57225 |
| **Eczema** | **Healthy Control** | Skin tissue | 28:9 | GPL14550 Agilent-028004 | GSE57225 |
| **Rheumatoid Arthritis** | **Healthy Control** | Synovial membrane | 33:20 | GPL96 Affymetrix U133A | GSE55457 |
| **Osteoporosis** | **Age matched control** | Bone marrow MSCs | 5:4 | GPL570 Affymetrix U133 Plus | GSE35958 |
| **Multiple Sclerosis** | **Healthy Control** | White blood cells | 8:5 | GPL6480 Agilent-014850 | GSE27688 |
| **SLE** | **Healthy Control** | Bone marrow endothelial progenitor cells | 6:6 | GPL11670 Affymetrix U133 Plus | GSE26951 |

**Table S3**. Comparison of Gene Ontology terms associated with Y102/Y202 cells and autoimmune disorders.

The ten most significantly enriched GO terms in Y02 samples and their significance in the disease datasets *p<0.05, **p<0.001, NS – Not significant (p>0.05).

| GO Term | p-value of enriched GO term | | | | | | |
|---|---|---|---|---|---|---|---|
| | Y02 | Pso | Ecz | RA | OP | MS | SLE |
| **Defense response** | 2.12E-17 | ** | ** | ** | NS | NS | NS |
| **Immune system process** | 5.96E-17 | ** | ** | ** | * | NS | NS |
| **Immune response** | 1.25E-14 | ** | ** | ** | NS | NS | NS |
| **Regulation of multicellular organismal process** | 1.07E-11 | ** | ** | NS | ** | NS | NS |
| **Response to external stimulus** | 5.48E-10 | ** | ** | ** | ** | NS | NS |
| **Regulation of cytokine production** | 6.54E-10 | ** | ** | NS | NS | NS | NS |
| **Positive regulation of multicellular organismal process** | 9.64E-10 | ** | ** | NS | * | NS | NS |
| **Regulation of immune effector process** | 1.05E-09 | ** | ** | ** | NS | NS | NS |
| **Response to other organism** | 1.84E-09 | ** | ** | ** | NS | NS | NS |
| **Response to stimulus** | 1.90E-09 | ** | ** | ** | ** | NS | NS |

**Table S4. Comparison of signalling pathways associated with Y102/Y202 cells and autoimmune disorders.**

The top ten most significantly upregulated pathways in Y02 were identified and compared with pathways significantly upregulated in the disease datasets *p<0.05, **p<0.001, NS – Not significant (p>0.05).

| Pathway Name | Y02 vs. Y01 | | p-value of up regulated pathway in disease samples | | | | | |
|---|---|---|---|---|---|---|---|---|
| | p value | Number of DEGs | Pso | Ecz | RA | OP | MS | SLE |
| Immunoregulatory interactions between a Lymphoid and a non-Lymphoid cell | 3.35E-10 | 30 | ** | ** | ** | NS | NS | NS |
| Interferon alpha-beta signaling | 8.78E-10 | 16 | ** | ** | NS | NS | NS | NS |
| GPCR downstream signaling | 4.60E-09 | 58 | ** | ** | ** | NS | NS | NS |
| GPCR ligand binding | 1.77E-08 | 53 | ** | ** | ** | NS | NS | NS |
| Type II interferon signaling | 3.04E-08 | 14 | ** | ** | ** | NS | NS | NS |
| Class I MHC mediated antigen processing and presentation | 4.31E-08 | 22 | NS | * | NS | NS | NS | NS |
| Allograft Rejection | 7.08E-08 | 21 | ** | ** | ** | NS | NS | NS |
| Integrin cell surface interactions | 2.42E-07 | 17 | * | ** | NS | ** | NS | ** |
| Gastrin-CREB signaling pathway via PKC and MAPK | 8.96E-06 | 24 | NS | ** | NS | NS | NS | NS |
| Cell surface interactions at the vascular wall | 1.07E-05 | 18 | ** | ** | ** | ** | NS | * |

**Table S5. Primer sequences**

| Target | Forward primer sequence | Reverse primer sequence |
|---|---|---|
| RPS27a | Tgg at gaga atg gca aaa tta gtc | CAC CCC AGC ACC ACA TTC A |
| CXCL10/IP10 | AAG CAG TTA GCA AGG AAA GGT CTA | GCA TCG ATT TTG CTC CCC TC |
| CXCL11 | CCT TGG CTG TGA TAT TGT GTG C | TGA ACA TGG GGA AGC CTT GAA |
| EPSTI1 | TGC ATA CAC CTT GAT AGC ACC AA | TCC TGC TCC GCA ATT CTT TG |
| HERC5 | GAG CTA AGA CCC TGT TTG G | CCA CCT TCC ACA TGC TAT C |
| IFI44L | TGC TCC TTC TGC CCC ATC TA | TGC TCC TTC TGC CCC ATC TA |
| ISG15 | ATG TCG GTG TCA GAG CTG AAG | GTT ATT CCT CAC CAG GAT GCT C |
| LY6E | CCT GGA GTC TTA CGG TCC AA | GTA CAC AGC CAG GCA CAC AT |
| MX1 | TTC AGC ACC TGA TGG CCT ATC | GTA CGT CTG GAG CAT GAA GAA CTG |
| MX2 | CAG AGG CAG CGG AAT CGT AA | TGA AGC TCT AGC TCG GTG TTC |
| RSAD2 | GTG GTT CCA GAA TTA TGG TGA GTA TTT | CCA CGG CCA ATA AGG ACA TT |
| SAA4 | GGA GAA AGG TCC ACA GCA CAA T | ATG TCC CAA TAG GCT CTG CC |
| CD54/ICAM1 | GCC AGG AGA CAC TGC AGA CA | TGG CTT CGT CAG AAT CAC GTT |
| CCL2/MCP1 | CAT AGC AGC CAC CTT CAT TCC | TCT CCT TGG CCA CAA TGG TC |
